# Collagen V Presented at the Cell-Substrate Interface Mimics Substrate Stiffening to Enhance Fibroblast Traction, Adhesion Turnover, and Persistent Migration

**DOI:** 10.64898/2026.10.21.760760

**Authors:** Shaina P. Royer, Himja Tiwari, David Cho, Sesha Sai Ramineni, Majid Rostami, Mohanish Chandurkar, Alexander X. Cartagena-Rivera, Sangyoon J. Han

## Abstract

Collagen type V (Col-V) is a minor fibrillar collagen that regulates collagen fibril assembly, and its haploinsufficiency causes classical Ehlers-Danlos syndrome, in which wound healing is delayed. Whether Col-V also changes how fibroblasts adhere to, pull on, and migrate across a collagen-presenting surface has not been tested directly. Here, we covalently coupled Col-I or Col-V to soft (2.7 kPa) silicone substrates and quantified the mechanical response of NIH/3T3 fibroblasts. On Col-V, cells generated about 1.5-fold higher traction, spread more, and formed smaller focal adhesions that each carried more force. Adhesions on Col-V assembled and disassembled faster. Single fibroblasts on Col-V migrated more slowly but more persistently, and fibroblast monolayers closed a 500 µm gap faster. Raising substrate stiffness to 13 kPa reproduced this phenotype on Col-I, including higher traction, larger spread area, and smaller focal adhesions, indicating that Col-V presentation produces an adhesion state that otherwise requires a stiffer substrate. In separate reconstituted collagen hydrogels, co-assembly with 10 to 20% Col-V yielded thinner, straighter fibrils, and 20% Col-V increased the indentation modulus of the gel. These results identify the collagen type presented at the cell-material interface as a design parameter that can partly substitute for bulk stiffness in controlling fibroblast traction and migration, and they suggest Col-V as a tunable component of collagen-based biomaterials.

**Statement of significance:** Substrate stiffness is a central design parameter for cell-instructive biomaterials, but it is difficult to change without also changing other material properties. We show that the type of collagen presented at the cell-material interface can stand in for a roughly fivefold increase in substrate stiffness. On soft substrates coupled with collagen V, a minor fibrillar collagen, fibroblasts generated traction, spreading, and compact focal adhesions comparable to those on a stiffer collagen I substrate, turned over adhesions faster, migrated more persistently, and closed wounds faster. Collagen V refined fibril architecture and stiffened reconstituted collagen hydrogels. These findings point to collagen V as a tunable component for collagen-based coatings and scaffolds intended to direct fibroblast migration, including wound repair.

## 1. Introduction

Classical Ehlers-Danlos syndrome (cEDS) is a heritable connective-tissue disorder caused in most patients by haploinsufficiency of collagen type V (Col-V), a quantitatively minor fibrillar collagen [1, 2] [3]. Col-V co-assembles with collagen I (Col-I) into heterotypic fibrils, where it regulates fibril nucleation and diameter, and its conservation from invertebrates to mammals points to a conserved role in matrix organization [4, 5]. Clinically, cEDS manifests as skin hyperextensibility, fragility, widened atrophic scars, and delayed wound healing [6]. cEDS skin exhibits reduced tensile strength and stiffness [7], even where fibril ultrastructure appears only mildly abnormal [8], suggesting that the consequences of Col-V deficiency are not limited to gross architectural defects.

In tissue, Col-V retains a partially processed N-terminal domain that projects from the fibril surface and is thought to limit lateral fibril growth [9, 10]. Loss of Col-V yields fewer, larger, and irregular fibrils and mechanically weaker connective tissue [8] [11]. Col-V deficiency also affects the cells that build and repair this matrix. Dermal fibroblasts from cEDS patients migrate poorly, a defect rescued by exogenous Col-V, and Col-V enhances fibroblast-mediated matrix contraction [12, 13]. Such effects could arise from differences in receptor engagement, in how the ligand is presented, or in matrix mechanics. Col-I and Col-V are recognized by an overlapping set of collagen-binding β1 integrins, and no integrin heterodimer specific to Col-V over Col-I has been identified (Supplementary Note 1) [14–21]. This raises the possibility that Col-V alters fibroblast behavior through the way it engages the mechanosensitive adhesion machinery of the cell rather than through a dedicated receptor [22].

Delayed wound healing is a hallmark of cEDS, and Col-V haploinsufficiency is associated with fragile dermis, impaired matrix remodeling, and altered dermal fibroblast behavior [23]. Wound closure requires fibroblasts to generate traction, maintain directional persistence, and advance collectively into the defect [24, 25], behaviors that depend on adhesion assembly, traction generation, and cytoskeletal remodeling [26]. These processes are strongly mechanoregulated. Fibroblasts probe their surroundings through integrin adhesions and adjust adhesion organization, traction, and motility to substrate stiffness, fibrillar architecture, and ligand presentation [27, 28], and soft substrates generally reduce traction and alter migration [29–31]. For the same reason, stiffness and ligand presentation are the two most widely used handles for engineering cell-instructive biomaterials [32, 33]. Whether the type of collagen presented to fibroblasts, independent of bulk stiffness, can shift their mechanical state has received little attention.

We previously proposed that Col-V acts as a mechanical modulator of the fibroblast microenvironment that tunes traction generation, migration persistence, and wound closure [34]. Here, we tested this idea at the cell-material interface. We covalently coupled Col-I or Col-V to soft silicone substrates, combined traction force microscopy, live-cell adhesion imaging, and single-cell and collective migration assays, and asked whether increasing substrate stiffness reproduces the response to Col-V. In parallel, we examined how Col-V co-assembly changes fibril architecture and stiffness in reconstituted collagen hydrogels. Our results show that Col-V presentation shifts fibroblasts into an adhesion and traction state that otherwise requires a stiffer substrate, and they identify Col-V as a candidate component for tuning collagen-based biomaterials.

## 2. Materials and methods

### 2.1. Cell culture

NIH/3T3 fibroblasts stably expressing paxillin-mRuby, a gift from the Ginsberg laboratory (University of California, San Diego) [35], were cultured in a 5% CO_2_, 37 °C incubator using Dulbecco’s Modified Eagle Medium (ThermoFisher) supplemented with 10% fetal calf serum, 1% penicillin/streptomycin, 1% non-essential amino acids, and 1% L-glutamine. The medium was not supplemented with ascorbic acid. For TFM and adhesion imaging, cells were seeded on coated substrates at approximately 20% confluency; TFM images were acquired 4 to 5 h after seeding, and adhesion dynamics were imaged starting 1.5 h after seeding. Single-cell migration imaging began 3 h after seeding. For the wound-closure assay, cells were cultured in the inserts for at least 24 h and confirmed to be confluent before the inserts were removed.

### 2.2. Substrate fabrication

High-refractive-index silicone gel substrates (Q-gel 920, CHT, Richmond, VA, USA) of defined stiffness were fabricated on glass-bottom dishes (#1 coverglass, MatTek Corporation) as previously described (Mittal et al., 2024). Gel stiffness was tuned by adjusting the mixing ratio of Q-gel 920 Part A to Part B; mixing ratios of 1:1.2 and 1:2 yielded substrate stiffnesses of 2.7 kPa and 13 kPa, respectively, as verified by measuring the shear storage modulus (G′) with a DHR-2 hybrid rheometer (TA Instruments, New Castle, DE, USA). Parts A and B were vortexed for 2 minutes and mixed by hand for 10 minutes, and 110 µL of the mixture was spin-coated onto a glass-bottom dish (250 rpm for 12 s with an acceleration of 100 rpm/s, then 1000 rpm for 30 s with an acceleration of 500 rpm/s) and cured in a convection oven at 80 °C for 2 hours, yielding a gel approximately 45 µm thick. Gels were stored in PBS for up to one week. Before bead and collagen coupling, the gel surface was silanized by immersion in 5% APTES for 3 minutes, rinsed three times with 96% ethanol, and air-dried.

### 2.3. Collagen coating protocol

Col-I or Col-V was diluted in PBS to a final concentration of 1 µg/mL in a total volume of 3.1 mL. Atelocollagen I (PureCol®, Advanced BioMatrix, Cat# 5005; bovine, pepsin-solubilized, approximately 3 mg/mL) was used for all Col-I conditions unless otherwise noted; results were compared against telocollagen I (Advanced BioMatrix) and showed no significant differences. Col-V was bovine type V collagen prepared by limited pepsin digestion and selective salt precipitation (SouthernBiotech, Cat# 1280-02S; 0.5 mg/mL in 0.5 M acetic acid); according to the manufacturer, purity is greater than 90% by SDS-PAGE, and we did not further characterize its chain composition. EDC [N-(3-dimethylaminopropyl)-N′-ethylcarbodiimide hydrochloride] was added to the collagen solution to a final concentration of 100 µg/mL to covalently couple collagen carboxyl groups to amine groups on an APTES (3-aminopropyltriethoxysilane)-silanized surface. The solution was briefly vortexed, poured onto the APTES-treated substrate, and incubated for 30 minutes at room temperature, followed by three rinses with PBS. The same protocol was used for silicone gels (TFM) and for glass-bottom dishes (adhesion imaging and single-cell migration). For the wound-closure assay, tissue culture plastic was plasma-treated and then coated with Col-I or Col-V at 1, 5, or 10 µg/mL without EDC.

### 2.4. Live-cell adhesion imaging

Fibroblasts expressing paxillin-mRuby were imaged using a total internal reflection fluorescence (TIRF) microscope (optoTIRF, CAIRN Research) mounted on a Nikon Ti-S microscope (Nikon Instruments) with a 60x objective. The microscope stage was equipped with an H301 stage-top incubator chamber and UNO controller (Okolab USA Inc, San Bruno, CA) to maintain cells at 5% CO_2_ and 37 °C in a humidified environment, and a CRISP autofocus unit (ASI Applied Scientific Instrumentation, USA) was used to maintain focus throughout imaging. Cells were seeded on 35 mm, No. 1.0 coverslip glass-bottom dishes (MatTek, USA) coupled with Col-I or Col-V as described above, and were imaged using a 587 nm laser with 300 ms exposure time every 15 seconds for 15 minutes, starting 1.5 h after seeding. Images were acquired with a Hamamatsu Orca-Flash 4.0 LT Plus CMOS camera (Hamamatsu Corporation, Bridgewater, NJ, USA) and controlled using MetaMorph imaging software (Molecular Devices, Downingtown, PA).

### 2.5. Focal adhesion analysis

Adhesion analysis was performed using the Focal Adhesion Package as previously described [36, 37]. Briefly, NAs were detected using a point-source detection method as previously described. FAs and FCs were segmented by thresholding, using a combination of Otsu and Rosin methods, on paxillin images processed with Gaussian filtering and background subtraction. Segmented areas larger than 0.2 µm² were classified as FCs (0.2–0.6 µm²) or FAs (≥0.6 µm²). The length and width of individual segmentations were evaluated as the major- and minor-axis lengths, respectively, of a fitted ellipse; eccentricity was calculated as the ratio of minor- to major-axis length. NAs were tracked over the full time-lapse sequence and compared against FC/FA segmentations to determine maturation state. Adhesion lifetimes were computed from the lifetimes of individual tracked trajectories, categorized by status (before adhesion, NA, FC, FA) at each time point.

### 2.6. Assembly and disassembly rate analysis

Assembly and disassembly rates were determined by measuring the background-subtracted fluorescence intensity of individual adhesions over time. Rate constants were determined from the slope of log(I/I₀) (assembly) and log(I₀/I) (disassembly) plots, where I₀ is the initial fluorescence intensity and I is the intensity at each subsequent time point [38]. To identify the most appropriate time window for rate determination, adjusted R² values were computed for multiple linear fits over different time ranges, and the fit with the highest adjusted R² was selected. Trajectories with an adjusted R² below 0.3 were excluded as noise.

### 2.7. Traction force imaging and analysis

Traction force microscopy (TFM) was performed on Q-gel 920 substrates as described above. Cells were cultured in Dulbecco’s Modified Eagle Medium without phenol red (Gibco, 31053) supplemented with 10% fetal bovine serum, 1% penicillin/streptomycin, 1% non-essential amino acids, and 1% L-glutamine. Prior to collagen coating, 40 nm carboxylate-modified dark-red fluorescent microspheres (FluoSpheres™, Invitrogen) were covalently conjugated to the gel surface via EDC-mediated coupling: 1 µL of beads and 5 µL of 100 mg/mL EDC were added to 5 mL of deionized water, sonicated for 15 minutes, poured onto the substrate, and incubated for 10 minutes at room temperature, followed by three PBS rinses and UV sterilization for 25 minutes, yielding a surface density of 1.3 bead/μm^2^. Live-cell imaging was performed on the same TIRF microscope setup described above, using 642 nm (beads) and 587 nm (paxillin) excitation with 300 ms exposure time, using the minimum illumination intensity needed for a clear image. Bead images of the relaxed gel were obtained after removing cells with 0.5 mL of 10% bleach. The displacement field was calculated from paired bead images (with and without cells) using a particle-tracking velocimetry method based on image cross-correlation, with a template size of 17 pixels and a maximum displacement of 50 pixels; outlier vectors were removed by filtering. Traction force reconstruction was performed using the Fast Boundary Element Method (FastBEM) with L2-norm-based regularization, with the regularization parameter selected via the L-curve corner method [39].

### 2.8. Single-cell migration assay

NIH/3T3 fibroblasts were cultured on glass substrates coupled with Col-I or Col-V (1 µg/mL) as described above and imaged by phase-contrast time-lapse microscopy for 24 hours at 10-minute intervals, beginning 3 h after seeding. Individual cell centroids were tracked using TrackMate at FIJI to generate migration trajectories. Mean speed, diffusion coefficient, and directional persistence (ratio of net to total path length) were calculated from the resulting trajectories using custom Matlab codes. Cumulative displacement distributions were computed from the maximum displacement of each tracked cell over the full 24-hour period.

### 2.9. Wound-closure (collective migration) assay

Fibroblast monolayers were cultured on tissue culture plastic coated with Col-I or Col-V, using a Culture-Insert 2 Well (ibidi) to define a 500 µm cell-free gap. Cells were cultured in the inserts for at least 24 h and confirmed to be confluent before the inserts were removed. The assay was first performed at 1 µg/mL, matching the coating density used for the single-cell migration experiments, and then repeated at 5 µg/mL, which produced a substantially larger separation between Col-I and Col-V; the main-text quantification (Fig. 4c–f) refers to the 5 µg/mL condition, and the 1 and 10 µg/mL data are shown in Supplementary Fig. S1. Upon removal of the insert, time-lapse phase-contrast imaging was performed with a 4× objective on an EVOS FL Auto imaging system (Smitha Rao laboratory, Michigan Technological University) at 2-hour intervals for up to 48 hours (1280 × 960 pixels; 1.83 µm/pixel). Wound boundaries were segmented in Fiji using the Wound Healing Size Tool macro [69], which applies contrast enhancement, a variance filter, thresholding, hole filling, and selection of the largest connected cell-free region; the variance window radius, threshold, and saturated-pixel percentage were held identical across all conditions and replicates. For each well the segmented open area was normalized to its own value at the first time point, so that every closure metric is expressed as a percentage of that well’s initial gap. Three metrics were derived per well. The wound-closure rate was obtained by linear regression over the pre-plateau portion of that well’s curve (normalized open area > 20%). The time to 50% closure (T₅₀) was obtained by fitting the full curve with a four-parameter logistic function and solving the fit for 50% of the initial open area; a straight-line fit was avoided because the post-closure plateau biases the fitted slope and therefore the extrapolated T₅₀. The time-averaged open area was obtained by trapezoidal integration of the normalized curve from 0 hr to a fixed endpoint, divided by that window; the endpoint was set per coating density to the longest interval common to every well at that density (46 hr at 5 and 10 µg/mL, 36 hr at 1 µg/mL), because an area integral is only comparable between wells when all are integrated over an identical interval. Residual open area at 24 hours, taken from the frame nearest 24 hours on each well’s own time base, is reported in Supplementary Fig. S2; it is not used in the main text because a single frame drawn from the steepest part of the Col-I curve is largely a noisier restatement of T₅₀. All metrics were computed per well rather than from a pooled average curve. Values lying more than 3 × the interquartile range beyond the first or third quartile were excluded before plotting and statistical testing, and Col-I and Col-V were compared with two-sided Wilcoxon rank-sum tests. At 5 µg/mL, N = 5 (Col-I) and N = 4 (Col-V) independent wells pooled from two independent experiments; at 1 µg/mL, N = 4 and N = 2; at 10 µg/mL, N = 1 per condition, which is reported descriptively without statistical testing.

### 2.10. Collagen gel SEM sample preparation and imaging

Collagen gels for SEM imaging were prepared for Col-I, 10% Col-V (90:10 Col-I:Col-V by mass), and 20% Col-V (80:20 Col-I:Col-V by mass) at a total collagen concentration of 1.5 mg/mL and a total volume of 100 µL (Col-I, bovine atelocollagen I, FibriCol®, Advanced BioMatrix, Cat# 5133, approximately 10 mg/mL; Col-V, SouthernBiotech, Cat# 1280-02S, 0.5 mg/mL). Atelocollagen I was used to match the pepsin-solubilized Col-V, and Col-V fractions above 20% were not used because of the low concentration of the Col-V stock. The required volume of Col-V was added to Col-I together with 10 µL of 10X PBS on ice, and deionized water was added to bring the Col-I and 10% Col-V mixtures to 70 µL before mixing. The pH was adjusted to 7.2 to 7.6 with 1 M and 0.1 M NaOH, deionized water was added as needed to a total volume of 100 µL, and the gels were polymerized at 37 °C for 90 minutes. Gels were fixed in 1% glutaraldehyde in 0.1 M phosphate buffer for 4 hours, rinsed three times for 5 minutes in deionized water, and dehydrated through a graded ethanol series (25%, 50%, 75%, 90%, and twice 100%; 10 minutes each). Samples were then rinsed twice for 10 minutes in hexamethyldisilazane (HMDS), placed on a glass coverslip, covered with HMDS, and left overnight for the HMDS to evaporate. Dried samples were sputter-coated with 5 nm of platinum/palladium at 20 mA. Imaging was performed on a Hitachi S-4700 cold field-emission scanning electron microscope (ACMAL, Michigan Technological University) at an accelerating voltage of 5 kV, an emission current of 10 µA, and a working distance of 12.5–12.7 mm. Ten images were acquired per gel at a nominal magnification of 20,000×, and the eight best images per condition were used for analysis. Because nominal magnification reflects the instrument display rather than the true imaged area, the actual horizontal field of view (≈6.3 µm) was calculated directly from the embedded scale bar of the acquired images. The same sample preparation and imaging protocol was used for all three collagen conditions.

### 2.11. Fiber orientation mapping

Orientation fields were computed using a custom MATLAB pipeline based on a steerable filter bank (2nd-order derivative of Gaussian), similar to previously published fibrillar alignment techniques [40]. Pixelwise orientation was assigned by identifying the maximal filter response across all angles, generating the color-coded orientation overlays. Fibril diameter was measured manually in ImageJ: starting from a corner of each image, the diameter of every clearly visible fibril was measured until 50 fibrils had been measured, and for fibrils of variable width the diameter was taken near the fibril midpoint, giving 400 fibrils per condition from eight images. For curvature, the steerable-filter response was followed by non-maximal suppression and segmentation of individual fibrils, and curvature was computed as the magnitude of the second derivative of each fibril centerline (units: 1/μm; N = 991 to 1259 fibrils per condition). Statistical comparisons were made using Kruskal–Wallis tests with Dunn–Šidák correction.

### 2.12. Collagen fibril sample preparation for atomic force microscopy

To prepare collagen fibrils for AFM imaging, freshly cleaved muscovite mica was used as the substrate. 72 µL of ECM buffer was added to the mica surface, followed by the collagen solution: 8 µL of Col-I (FibriCol®, Advanced BioMatrix, Cat# 5133, diluted to 1 mg/mL) for 100% Col-I samples, 8 µL of Col-V (0.5 mg/mL; SouthernBiotech, Cat# 1280-02S) for 100% Col-V samples, or 6 µL of Col-I (1 mg/mL) plus 2 µL of Col-V (0.5 mg/mL) for 80:20 Col-I/Col-V blend samples, each gently mixed into the ECM buffer droplet by pipetting without touching the mica surface. The mica disc was placed in a 60 mm petri dish with a wet Kimwipe, sealed with parafilm, and incubated overnight at room temperature. Samples were then rinsed five times with 1X PBS and covered with fresh ECM buffer.

### 2.13. Collagen hydrogel sample preparation for atomic force microscopy

Collagen hydrogels for bulk AFM microindentations were prepared on 35 mm glass-bottom dishes (MatTek Corporation) following a standard collagen gel polymerization protocol [41] (10X PBS, deionized water, and acid-solubilized collagen stock solution mixed on ice in that order, with pH adjusted to the physiological range using NaOH), adapted for two collagen conditions as follows. For 100% Col-I hydrogels, 30 µL of 10X PBS was mixed with 45 µL of Col-I stock solution (FibriCol®, Advanced BioMatrix, Cat# 5133; approximately 10 mg/mL). For 80:20 Col-I/Col-V blend hydrogels, 30 µL of 10X PBS was mixed with 36 µL of Col-I stock solution and 180 µL of Col-V stock solution (0.5 mg/mL; SouthernBiotech, Cat# 1280-02S). The mixture was titrated to pH 7.2–7.6 with 1 M and 0.1 M NaOH and brought to a total volume of 300 µL with deionized water. 150 µL of the resulting solution was deposited onto a MatTek dish and spread to coat the entire glass surface, then incubated for 2 hours at 37 °C to allow hydrogel polymerization. Assuming uniform spreading over the 20 mm glass well, the resulting hydrogels were approximately 0.5 mm thick.

### 2.14. Atomic force microscopy

Atomic force microscopy was performed using a JPK NanoWizard ULTRA Speed 2 BioAFM system (Bruker) mounted on a Ti2-E inverted microscope (Nikon) equipped with a CSU-W1 spinning disk confocal unit (Yokogawa) and an ORCA-Flash 4.0 V3 digital CMOS camera (Hamamatsu). Two complementary AFM approaches were used: quantitative imaging (QI) mode for collagen fibrils on mica, using ultra-sharp PEAKFORCE-HIRS-F-B probes (Bruker); and single-point force spectroscopy for bulk collagen hydrogels, using soft silicon nitride cantilevers with an attached 45 µm diameter polystyrene spherical probe (Novascan).

For QI imaging, probes were calibrated at room temperature in PBS using the instrument’s built-in contact-based calibration method (cantilever spring constants: 0.045–0.05 N/m) and then thermally equilibrated to 37 °C for 30 minutes. Collagen fibrils on mica were imaged using a force setpoint of 50–250 pN, a Z-length of approximately 250 nm, and a Z-speed of 25 µm/s, at a resolution of 512 × 512 pixels, with multiple images collected per specimen across triplicate experiments. Fibril thickness was quantified from line profiles traced across individual fibrils using JPKSPM Data Processing software.

For force spectroscopy on collagen hydrogels, probes were calibrated as above (cantilever spring constant: 0.71 N/m). Force curves were recorded using a 5 nN force setpoint, a 2500 nm Z-length, and a 2.5 µm/s Z-speed. At least five replicate force curves were collected per location, with approximately 10 seconds between measurements to allow hydrogel recovery, and at least five locations were probed per sample across triplicate experiments per condition. Young’s modulus was extracted using JPKSPM Data Processing software, fitting the Hertz contact model for a spherical probe to each force curve up to 1 nN [42, 43].

### 2.15. Statistics

Statistical analyses were performed using the MATLAB Statistics Toolbox. Two-group comparisons were performed with the two-sided Wilcoxon rank-sum test, which is equivalent to the Mann-Whitney U test. The three conditions in Fig. 7 were compared using one-way ANOVA with Tukey’s post-hoc test, and fibril diameter and curvature (Fig. 5) were compared using the Kruskal-Wallis test with Dunn-Šidák correction. The proportion of cells exceeding a displacement threshold (Fig. 3g) was compared using a two-sided Fisher’s exact test. Unless stated otherwise, significance is indicated as *P < 0.05, **P < 10⁻³, ***P < 10⁻⁵, and ****P < 10⁻⁸**. Values more than 3 × the interquartile range beyond the first or third quartile were excluded before plotting and testing. The number of cells, adhesions, wells, or measurements for each experiment is reported in the corresponding figure legend**

## 3. Results

### 3.1. Col-V increases fibroblast traction and shifts focal adhesion organization on soft substrates

To test whether Col-V changes the mechanical response of fibroblasts, we performed traction force microscopy (TFM) on NIH/3T3 fibroblasts plated on soft (2.7 kPa) silicone substrates to which bovine atelocollagen I (Col-I; PureCol®, Advanced BioMatrix) or bovine Col-V (SouthernBiotech), both pepsin-solubilized, was covalently coupled from solutions of the same concentration (1 µg/mL; Section 2.3). Because the coating is a thin molecular layer on a much thicker elastomer, the bulk stiffness of the substrate is set by the silicone and is identical for both coatings, so differences between conditions reflect the collagen presented at the interface. Cells were imaged within 5 h of seeding, which limits any contribution from cell-deposited matrix. Fluorescent beads (40 nm) on the gel surface enabled simultaneous measurement of cell morphology and substrate deformation (**Fig. 1b**). Fibroblasts on Col-V displayed more polarized morphologies, larger substrate deformation (**Fig. 1c**), and higher traction (**Fig. 1d**) than those on Col-I. Mean displacement per cell increased about twofold (**Fig. 1e**) and mean traction about 1.5-fold (**Fig. 1f**) on Col-V. Cell spread area, another stiffness-sensitive phenotype [32, 33], was also larger on Col-V (**Fig. 1g**). To examine adhesion organization, cells expressing paxillin-mRuby were imaged under identical conditions (**Fig. 1h**) and analyzed with our Focal Adhesion Package [37] (**Fig. 1i**). Focal adhesions (FAs) were smaller on Col-V (**Fig. 1j**), and FA density per cell area was lower across the whole cell, the cell periphery, and the cell interior, with the largest reduction in the interior (**Fig. 1k**). In contrast, nascent adhesion (NA) density was more than twofold higher on Col-V (**Fig. 1l**), indicating that Col-V does not suppress adhesion formation but shifts the adhesion population toward small, early-stage contacts. Traction normalized to the number of FAs was higher on Col-V (**Fig. 1m**), so each mature adhesion carried more force despite its smaller size. Because cells on Col-V also spread more, the lower FA density reflects mature adhesions distributed over a larger footprint rather than a loss of adhesive capacity. Thus, on substrates of identical stiffness, Col-V presentation increases traction while favoring numerous small adhesions that each bear more load.

**Figure 1.**
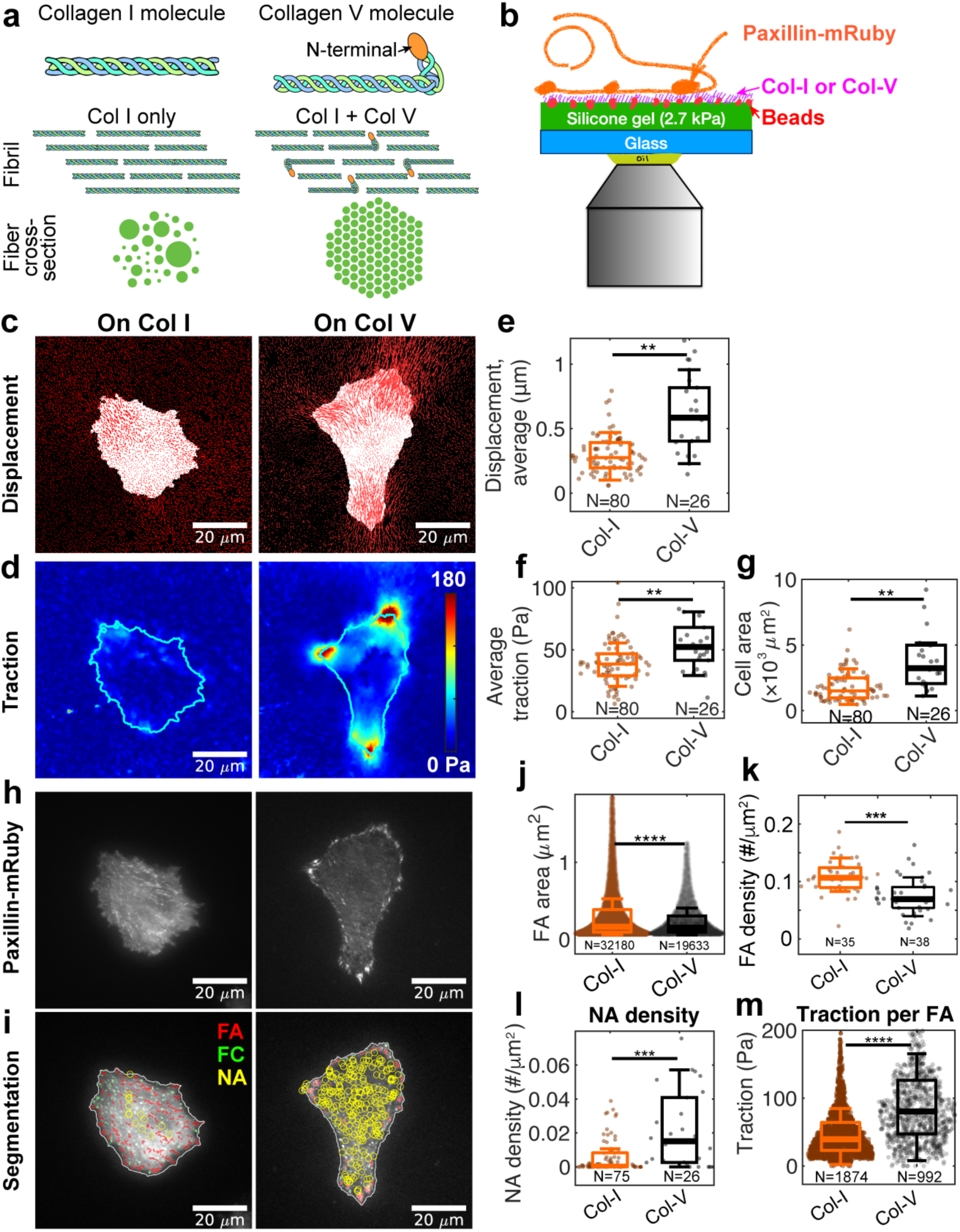
Col-V increases fibroblast traction and shifts focal adhesion organization on soft substrates. **(a)** Schematic of Col-I and Col-V molecules and, for reference, their co-assembly into heterotypic fibrils in tissue, where Col-V retains a partially processed N-terminal domain. In this study, both collagens were presented as molecules covalently coupled to the substrate (Section 2.3), and the pepsin-extracted Col-V used here is likely to lack part of this domain. **(b)** Schematic of the traction force microscopy (TFM) setup. Paxillin-mRuby-expressing NIH/3T3 fibroblasts were plated on 2.7 kPa silicone gels coupled with Col-I or Col-V (1 µg/mL), whose surface carries 40 nm fluorescent beads for measuring cell-generated substrate deformation. **(c)** Representative bead displacement fields for fibroblasts on Col-I and Col-V. Displacement vectors are shown in red overlaid on cell masks (white). **(d)** Corresponding traction-stress maps reconstructed from bead displacements (Section 2.7). Cells on Col-V display higher traction magnitudes and broader stress distributions than those on Col-I. **(e–g)** Quantification of mean bead displacement (e), mean traction stress (f), and projected cell area (g) per cell. **(h)** Representative paxillin-mRuby images showing focal adhesions (FAs) on Col-I and Col-V substrates. **(i)** Segmentation maps of adhesions categorized as nascent adhesions (NAs, yellow), focal complexes (FCs, green), and FAs (red). **(j–l)** Quantification of FA area (j), FA density (k), and NA density (l). FAs were smaller and less dense on Col-V across the whole cell, periphery, and interior, whereas NA density was more than twofold higher. **(m)** Mean traction normalized to the number of FAs per cell. Scale bars, 20 µm. Statistical significance: two-sided Wilcoxon rank-sum test; *P < 0.05, **P < 10⁻³, ***P < 10⁻⁵, ****P < 10⁻⁸.

### 3.2. Col-V accelerates focal adhesion assembly and disassembly

Because adhesions on Col-V were smaller yet carried more force (**Fig. 1j–m**), we asked whether Col-V also changes adhesion kinetics, which directly influence traction transmission and migration persistence [44, 45]. Using our FA tracking software [36], we segmented and tracked thousands of adhesion trajectories in paxillin-mRuby-expressing NIH/3T3 fibroblasts on glass coupled with Col-I or Col-V (**Supplementary Videos 1 and 2**). These experiments were performed on glass to optimize TIRF imaging, so they isolate the effect of the presented collagen on a stiff substrate. Color-coding individual trajectories by assembly rate highlighted faster assembly on Col-V, primarily near the cell edge (**Fig. 2a,b**). Zoomed time series of representative adhesions showed steeper rises in paxillin intensity on Col-V (**Fig. 2c,d**), and their log-normalized intensity trajectories built up faster (**Fig. 2e**). The apparent assembly rate constant [38] across all tracked adhesions was higher on Col-V (**Fig. 2f**). In contrast, the nucleation rate of NAs at the protrusive edge was lower on Col-V (**Fig. 2g**). Adhesions on Col-V also disassembled faster. Time-lapse sequences of adhesions entering the turnover phase (**Fig. 2h,i**) and the corresponding log-normalized inverted intensity traces (**Fig. 2j**) showed steeper decay, and the apparent disassembly rate constant was significantly higher on Col-V (**Fig. 2k**). Thus, on Col-V, fewer adhesions nucleate per unit time at the edge, but those that form grow and turn over faster. Together with the compact, highly loaded adhesions on soft Col-V substrates (**Fig. 1**), these data suggest that Col-V shifts adhesions toward a rapidly cycling, highly loaded state rather than toward slow, size-based maturation.

**Figure 2.**
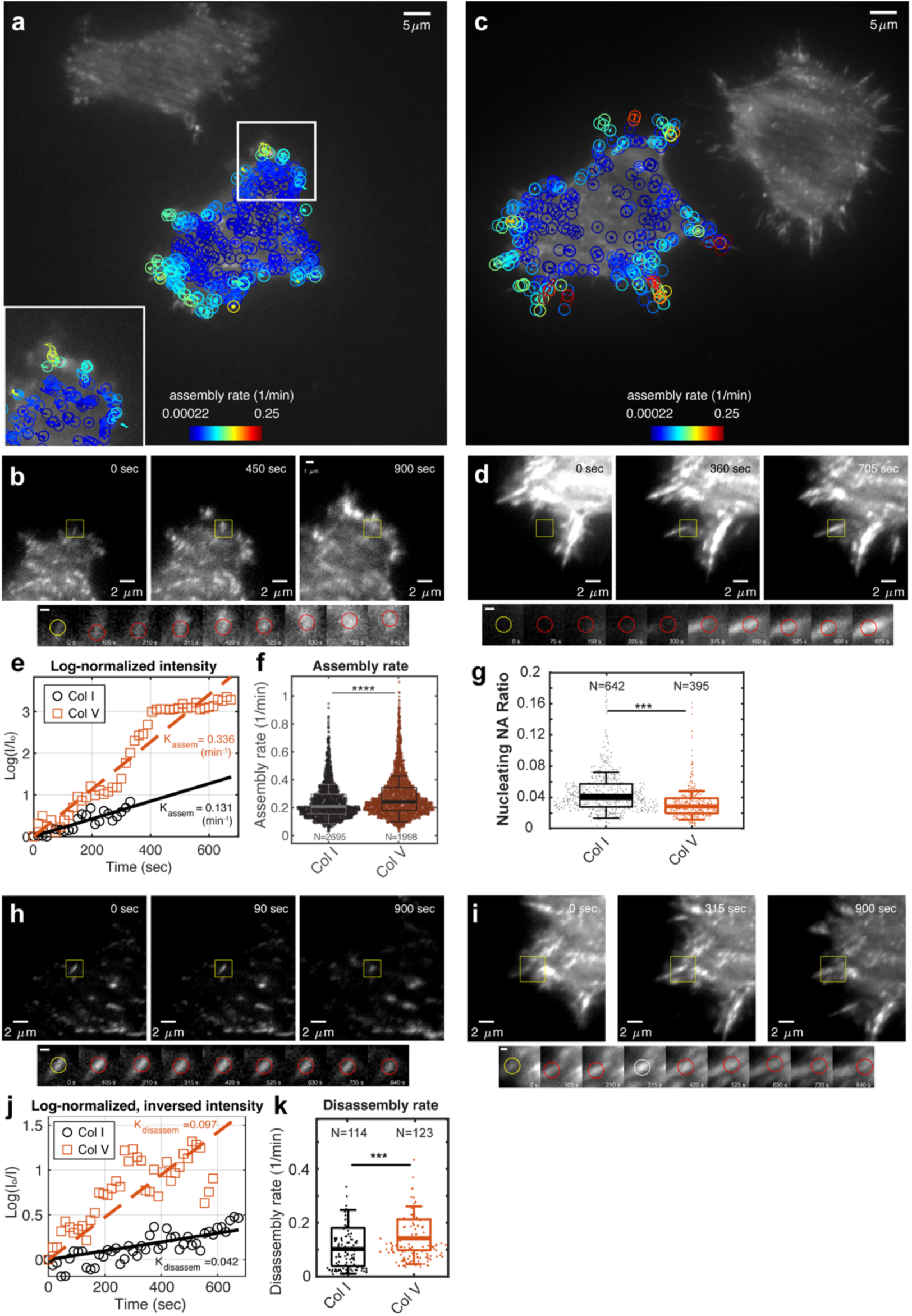
Col-V accelerates focal adhesion assembly and disassembly. Paxillin-mRuby-expressing NIH/3T3 fibroblasts on glass coupled with Col-I or Col-V (1 µg/mL). **(a–b)** Representative paxillin-mRuby images of fibroblasts on Col-I (a) or Col-V (b), color-coded by local assembly rate. Adhesions on Col-V show elevated assembly rates, particularly at the protrusive edge. **(c–d)** Zoomed time-lapse views of boxed adhesions from (a–b) illustrating faster paxillin intensity increase on Col-V. Insets show tracked regions with corresponding kymographs. **(e)** Population-averaged, log-normalized intensity trajectories of representative tracked adhesions reveal a steeper rise on Col-V. **(f)** Distribution of single-adhesion assembly rates quantified from intensity slopes, showing higher rates on Col-V. **(g)** Nucleation rate of NAs at the protrusive edge, which was lower on Col-V. **(h–i)** Time-lapse sequences of adhesions undergoing disassembly on Col-I (h) and Col-V (i); boxed ROIs correspond to tracked adhesions shown in lower kymographs. **(j)** Log-normalized, inverted intensity traces of representative adhesions show faster decay on Col-V than on Col-I. **(k)** Distribution of single-adhesion disassembly rates, showing faster disassembly on Col-V. Statistical significance: two-sided Wilcoxon rank-sum test; *P < 0.05, **P < 10⁻³, ***P < 10⁻⁵, ****P < 10⁻⁸**; error bars, s.e.m. Scale bars: 5 µm (a–b); 2 µm (c–d, h–i). See Supplementary Videos 1 and 2 for representative adhesion tracking**

### 3.3. Col-V enhances migration persistence and effective displacement of fibroblasts

We next asked whether these differences in traction and adhesion turnover alter single-cell migration. NIH/3T3 fibroblasts were cultured on glass coupled with Col-I or Col-V (1 µg/mL), and phase-contrast time-lapse imaging was performed for 24 hours. Individual cell centroids were tracked to generate migration trajectories, from which motility parameters were extracted. Representative trajectories (**Fig. 3a,b**) revealed distinct behaviors: cells on Col-I followed broader but more erratic paths, whereas those on Col-V followed straighter trajectories with smaller instantaneous displacements. Mean migration speed was slightly lower on Col-V than on Col-I (**Fig. 3c**), and the diffusion coefficient, which reflects the overall spread of cell positions, was also lower (**Fig. 3d**). In contrast, directional persistence, the ratio of net to total path length, was significantly higher on Col-V (**Fig. 3e**). This trade-off between lower speed and higher persistence resulted in greater effective displacement. Cumulative displacement distributions (**Fig. 3f**) showed that a larger fraction of cells on Col-V exceeded a 150 µm displacement threshold than on Col-I (**Fig. 3g**; N = 91 Col-I and N = 33 Col-V cells; two-sided Fisher’s exact test), despite their lower speed. Thus, Col-V promotes directionally persistent migration that allows fibroblasts to cover longer distances despite moving more slowly, consistent with the faster adhesion turnover in **Fig. 2**.

**Figure 3.**
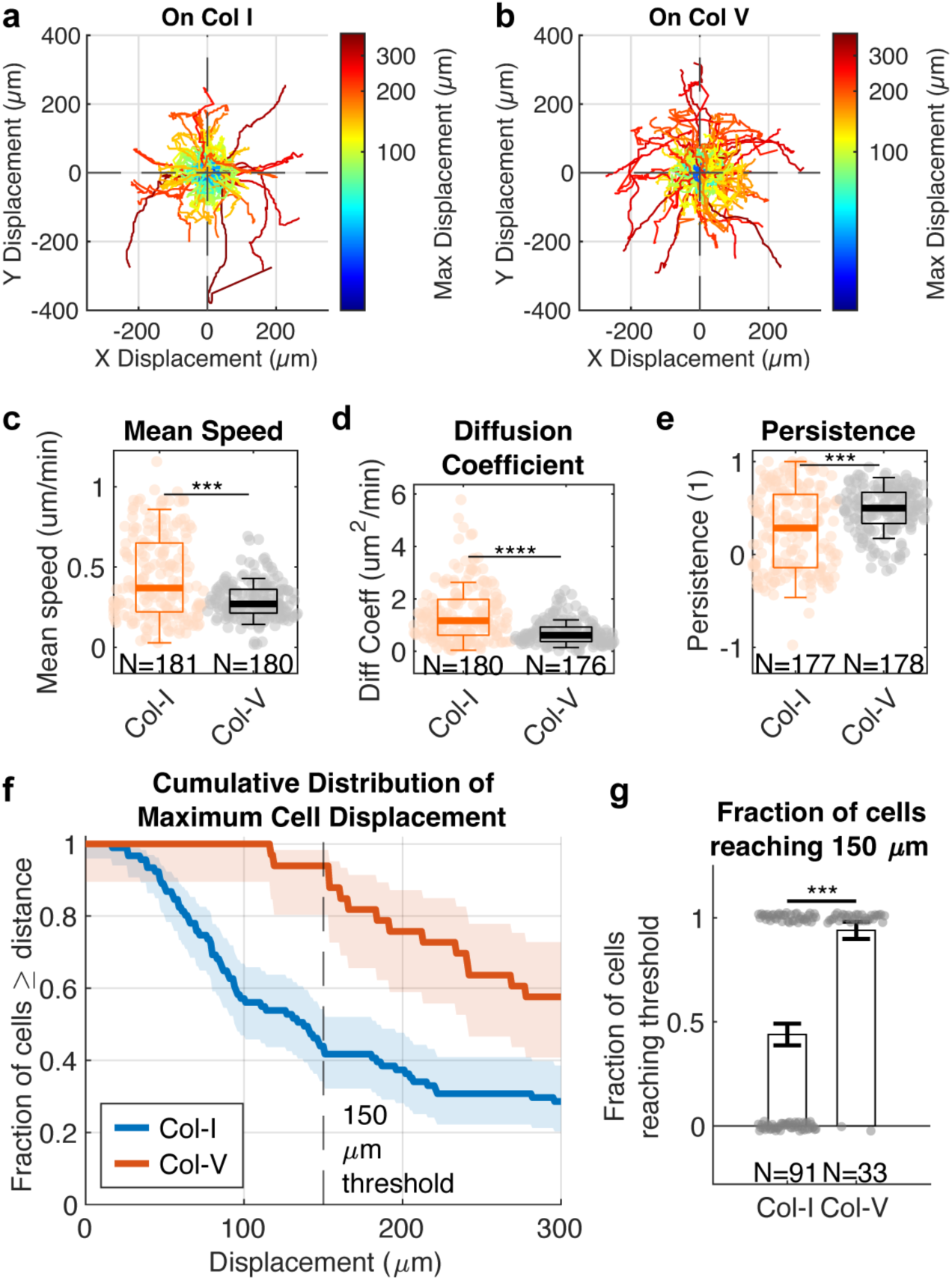
Col-V increases fibroblast migration persistence and effective displacement despite reduced speed. **(a–b)** Representative migration trajectories of NIH/3T3 fibroblasts on glass coupled with Col-I (a) or Col-V (b), color-coded by maximum cell displacement. Each line represents the centroid path of an individual cell tracked for 24 hours. **(c–e)** Mean cell speed (c), diffusion coefficient (d), and directional persistence, the ratio of net to total path length (e). **(f)** Cumulative distribution of maximum cell displacement over 24 hours. Shaded bands indicate 95% Wilson score confidence intervals for the proportion of cells exceeding each displacement value. **(g)** Fraction of cells exceeding a 150 µm displacement threshold. Gray points show the binary outcome (reached or did not reach threshold) for each cell, jittered for visibility; bars show the proportion per condition, and error bars are binomial s.e.m. (N = 91 Col-I, N = 33 Col-V). Statistical significance: two-sided Wilcoxon rank-sum test for (c–e) and two-sided Fisher’s exact test for (g); *P < 0.05, **P < 10⁻³, ***P < 10⁻⁵, ****P < 10⁻⁸**; error bars, s.e.m. except in (g**

### 3.4. Col-V accelerates fibroblast collective migration and wound closure

To determine whether Col-V enhances collective fibroblast migration at the multicellular level, we performed an *in-vitro* wound-closure assay using a Culture-Insert 2 Well (ibidi) on tissue culture plastic coated with either pure Col-I or Col-V. At 1 µg/mL, the coating density used for the single-cell experiments, the closure rate, T₅₀ and integrated open area all favoured Col-V on average, but the distributions overlapped and no comparison approached significance (**Supplementary Fig. S1**; N = 4 Col-I and N = 2 Col-V, a design for which the smallest attainable two-sided rank-sum P value is 0.13, so this comparison is under-powered by construction rather than negative). Because the separation between substrates appeared to grow with the amount of collagen presented to the cells, we repeated the assay at 5 µg/mL, where Col-I and Col-V separated clearly; all quantification reported below refers to that coating density. Upon removal of the insert, fibroblast monolayers initiated directed migration into the cell-free region (**Fig. 4a**). Time-lapse imaging for 46–78 hr revealed markedly faster closure on Col-V than on Col-I (**Fig. 4b**, **Supplementary Videos 3 and 4**). Normalized open-area curves declined more steeply on Col-V, separating from Col-I within the first ∼8 hr and remaining separated for the rest of the ∼46 hr imaging window (**Fig. 4c**). Fitting the pre-plateau portion of each replicate’s curve revealed a significantly higher wound-closure rate on Col-V than on Col-I (3.00 ± 0.91 vs. 2.15 ± 0.22 % of the initial open area per hour, mean ± SD; P = 0.016; **Fig. 4d**). Consistent with this, the time required to reach 50% closure (T₅₀), obtained from a logistic fit to each replicate’s own closure curve, was significantly shorter on Col-V (16.7 ± 3.4 vs. 23.6 ± 2.2 hr; P = 0.016; **Fig. 4e**). Because both of these measures summarize the curve by a single fitted parameter, we also integrated each well’s closure curve over the first 46 hr, the longest interval common to every well at this coating density. The resulting time-averaged open area was likewise significantly lower on Col-V (37.1 ± 6.8 vs. 51.0 ± 4.0 % of the initial gap; P = 0.016; **Fig. 4f**), showing that the advantage holds across the entire closure trajectory and not only at the fitted midpoint. The residual open area at a single 24 hr timepoint trended in the same direction but did not reach significance at this replicate number (**Supplementary Fig. S2**). Data are from N = 5 (Col-I) and N = 4 (Col-V) independent wells pooled from two independent experiments. Together, these data demonstrate that Col-V not only enhances single-cell traction, adhesion turnover, and migration persistence (**Figs. 1–3**) but also accelerates multicellular wound closure, consistent with, although not a direct measure of, more persistent collective migration.

**Figure 4.**
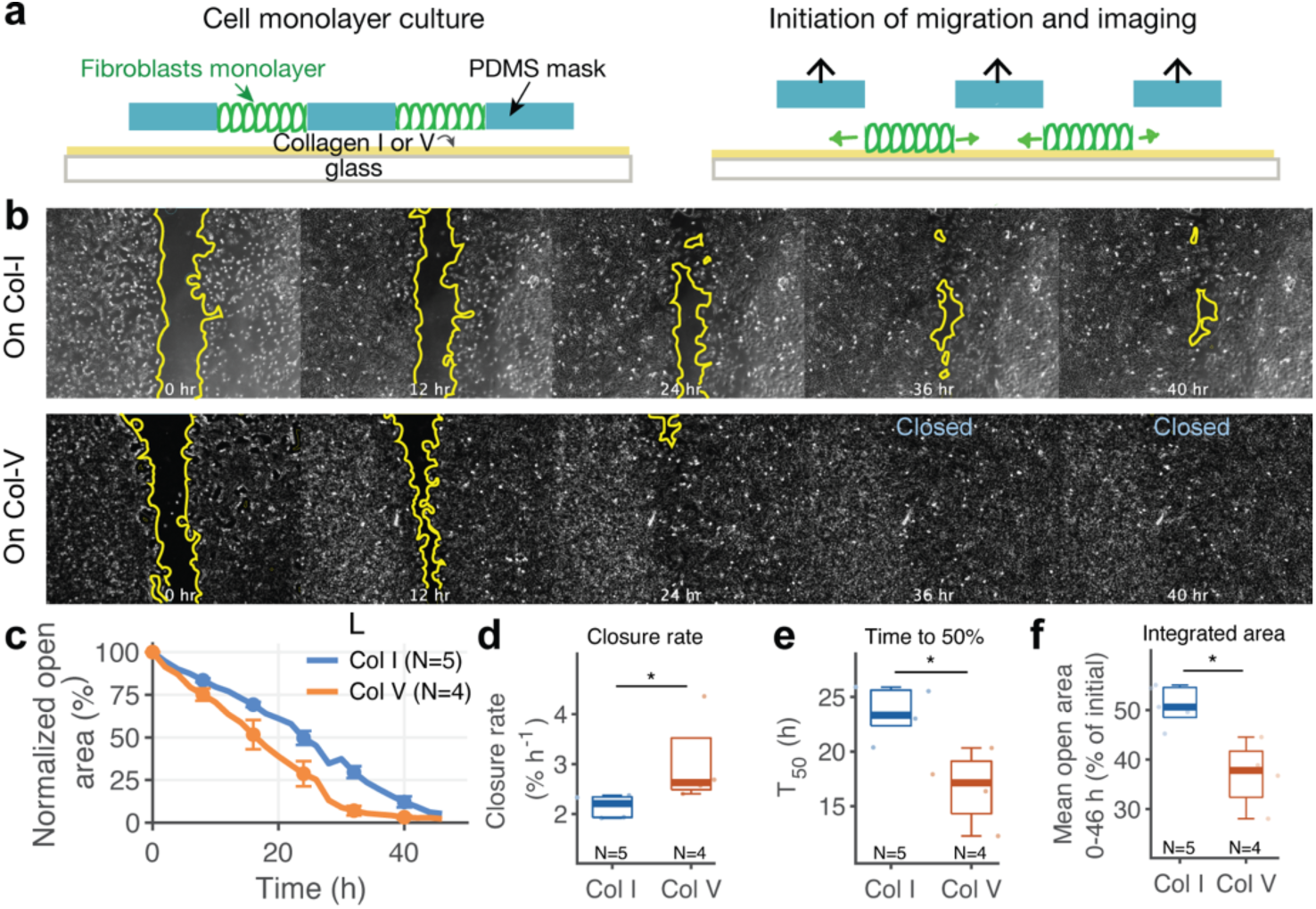
Col-V accelerates wound healing in vitro. Fibroblast monolayers on tissue culture plastic coated with 5 µg/mL Col-I or Col-V. **(a)** Schematic of the wound-closure assay using a Culture-Insert 2 Well (ibidi), which defines a 500 µm cell-free gap on the collagen-coated surface. Upon removal of the insert, fibroblasts migrate inward toward the gap. **(b)** Representative time-lapse images of wound closure on Col-I (top) and Col-V (bottom) at the indicated times. Yellow outlines denote wound boundaries segmented in Fiji. **(c)** Normalized open-area curves (mean ± SEM), each well normalized to its own open area at t = 0; error bars are plotted every 8 hr for clarity. Curves are shown over the interval in which every replicate of that condition still contributes data (0–46 hr). **(d)** Wound-closure rate, from linear regression over the pre-plateau portion of each replicate’s curve (normalized open area > 20%). **(e)** Time to 50% closure (T₅₀), obtained by fitting each replicate’s full curve with a four-parameter logistic function and solving it for 50% of that well’s initial open area. **(f)** Time-averaged open area, obtained by integrating each well’s normalized closure curve from 0 to 46 hr, the longest interval common to every well at this coating density, and dividing by that window. Unlike (d) and (e), this measure uses the whole trajectory rather than a single fitted parameter. In (d–f), boxes span the interquartile range with the median marked, whiskers extend to the most extreme values lying within 0.5 × the interquartile range of the box, and the overlaid symbols are individual wells; outliers are not drawn as separate symbols. Values more than 3 × the interquartile range beyond the first or third quartile were excluded before plotting and testing. N = 5 (Col-I) and N = 4 (Col-V) independent wells pooled from two independent experiments. Significance is indicated as *, P < 0.05; **, P < 10⁻³; ***, P < 10⁻⁵; ****, P < 10⁻⁸; two-sided Wilcoxon rank-sum test. With N = 5 and N = 4 the smallest attainable two-sided rank-sum P value is 0.016, which is reached in all three of (d), (e) and (f). The residual open area at 24 hr is shown in Supplementary Fig. S2, and the corresponding measurements at 1 and 10 µg/mL in Supplementary Fig. S1.

### 3.5. Col-V co-assembly yields thinner and straighter fibrils in reconstituted collagen hydrogels

The experiments above used thin molecular coatings on substrates of fixed stiffness. In tissue and in collagen-based scaffolds, however, Col-V acts as a co-assembling component of heterotypic Col-I/Col-V fibrils, where it regulates fibril nucleation and diameter [46, 47] [10]. To ask how Col-V changes the structure and mechanics of a collagen material in this setting, we characterized reconstituted Col-I hydrogels with and without Col-V. These experiments address a different length scale from the coatings and are presented as a parallel characterization of Col-V as a matrix component, not as a description of the coated surfaces. We imaged by SEM hydrogels polymerized at 1.5 mg/mL total collagen from Col-I alone or with 10% or 20% Col-V by mass. Pure Col-I gels displayed thick, coarse fibrils that were loosely organized and heterogeneous in diameter (**Fig. 5a**). In contrast, networks containing Col-V exhibited visibly finer fibrils and a denser meshwork (**Fig. 5b,c**). Automated fiber-orientation mapping showed that Col-I networks were broadly distributed in orientation with irregular local waviness (**Fig. 5d**), whereas networks containing 10% or 20% Col-V showed tighter local alignment and more uniform directional coherence (**Fig. 5e,f**).

**Figure 5.**
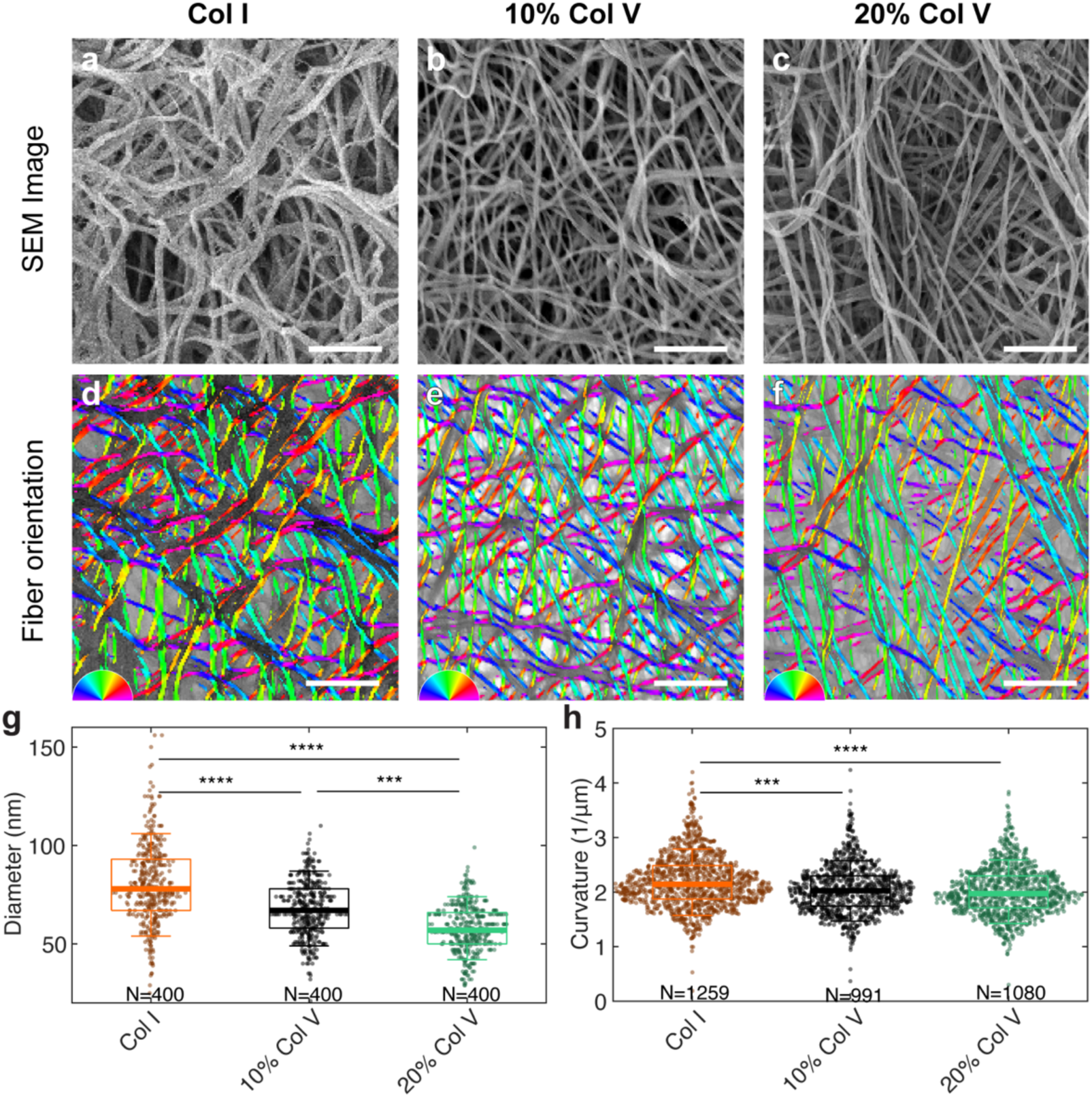
Col-V incorporation refines fibril architecture in reconstituted collagen networks. **a–c**, Representative SEM images of 1.5 mg/mL collagen gels polymerized with **Col-I only**, **10% Col-V**, or **20% Col-V**. Col-I networks contain thick, coarse fibrils, whereas Col-V incorporation produces visibly thinner and denser fibrillar meshes. Scale bars, 2 µm. **d–f**, Automated fiber-orientation maps from the same fields, with color indicating local fibril direction. Col-V increases local alignment and reduces long-wavelength waviness. **g**, Fiber-diameter quantification (N = 400 fibrils per condition). Col-V significantly reduces fibril thickness. **h**, Fibril-curvature analysis (N = 1259, 991, 1080 fibers for Col-I, 10% Col-V, 20% Col-V). Addition of Col-V decreases curvature, indicating straighter fibrils. Statistical significance: Kruskal-Wallis test with Dunn-Šidák correction; ***P < 0.001; ****P < 0.0001.

Quantitative analysis supported these observations. Fibril diameter decreased with Col-V content, with the largest diameters in Col-I alone, followed by 10% and then 20% Col-V (**Fig. 5g**). Fibril curvature, a measure of local tortuosity, was also lower in Col-V-containing networks (**Fig. 5h**), indicating straighter, less undulating fibrils. Thinner fibrils are consistent with the established role of Col-V in limiting lateral fibril growth *[46, 47]* [10], whereas reduced curvature was not anticipated and suggests that Col-V also changes how fibrils are organized within the network. Because SEM requires dehydration, absolute diameters are likely smaller than in the hydrated state, but all conditions were processed identically, so relative differences can be compared.

### 3.6. AFM reveals distinct Col-V assembly on mica and stiffening of Col-V-containing hydrogels

We next used atomic force microscopy (AFM). To visualize the assembly products of each collagen at the nanoscale, we first imaged fibrils formed from dilute collagen solutions on mica (**Fig. 6a**). Col-I formed long fibrils with strong uniaxial alignment (**Fig. 6a**, top), consistent with the known tendency of collagen to orient on atomically flat, negatively charged surfaces [48–50]. Col-V instead formed dense, irregular, three-dimensional aggregates with short, multidirectional protrusions (**Fig. 6a**, bottom), resembling morphologies attributed to Col-V-driven fibril nucleation [5, 47]. These mica preparations differ from the hydrogels in **Fig. 5** in substrate, concentration, and composition, so we use them only as a qualitative illustration of the distinct self-assembly behavior of Col-V. The high-aspect-ratio topography of the Col-V aggregates also precluded reliable indentation of individual fibrils [42, 51], so we did not attempt to derive fibril-level moduli.

**Figure 6.**
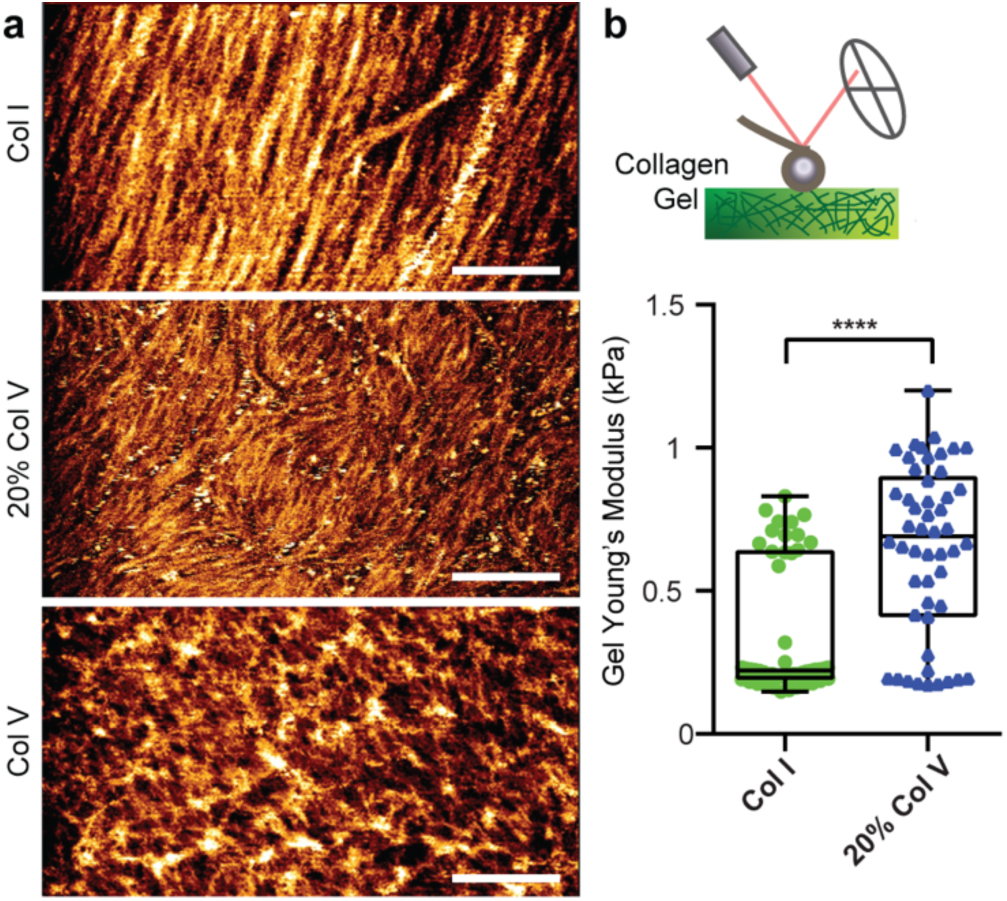
AFM imaging of collagen assemblies on mica and indentation of collagen hydrogels. **(a)** Representative AFM height images of collagen assemblies formed on mica from Col-I (1 mg/mL stock) or Col-V (0.5 mg/mL stock). Col-I forms long, aligned fibrils on the negatively charged mica surface, whereas Col-V forms short, irregular, highly protrusive aggregates. These images illustrate self-assembly behavior and were not used for mechanical measurements. **(b)** Apparent Young’s modulus of 1.5 mg/mL Col-I and 80:20 Col-I/Col-V hydrogels measured by spherical-probe AFM indentation on the hydrated gel surface. Col-V-containing hydrogels are significantly stiffer than Col-I hydrogels. Box plots show median, interquartile range, and all data points (N = 50 force curves per condition). ****P < 0.0001.

To measure network-level mechanics, we performed AFM microindentation with a 45 µm spherical probe on hydrated 1.5 mg/mL collagen hydrogels (**Fig. 6b**, top). The gel surfaces were too rough for high-resolution imaging but suitable for force spectroscopy. Hydrogels containing 20% Col-V had a significantly higher apparent Young’s modulus than Col-I hydrogels (**Fig. 6b**, bottom). The hydrogels were approximately 0.5 mm thick (150 µL spread over a 20 mm glass well), whereas the indentation depth within the fitted force range was below 1 µm, so substrate effects on the fitted modulus are expected to be negligible [42, 43]. Thus, although Col-V produced thinner fibrils, it increased the stiffness of the network as a whole, an apparent paradox that we discuss below in terms of fibril straightness.

### 3.7. Increasing substrate stiffness reproduces the Col-V phenotype on Col-I

On soft substrates of fixed stiffness, Col-V presentation raised traction and spread area and produced small, highly loaded adhesions. Because increasing substrate stiffness typically raises traction and spreading [29–31], we asked whether a stiffer Col-I substrate reproduces the Col-V phenotype. Fibroblasts were cultured on 13 kPa silicone substrates coupled with Col-I and compared with cells on 2.7 kPa substrates coupled with Col-I or Col-V. Mean bead displacement per cell was similar on Col-I/2.7 kPa and Col-I/13 kPa and larger on Col-V/2.7 kPa (**Fig. 7a,b**). Because traction depends on both displacement and substrate stiffness, similar displacements on a stiffer substrate imply larger forces. Traction reconstruction confirmed this: mean traction was significantly higher on Col-I/13 kPa than on Col-I/2.7 kPa and reached the level measured on Col-V/2.7 kPa (**Fig. 7c,d**). Cell spread area was likewise larger on Col-I/13 kPa than on Col-I/2.7 kPa and similar to that on Col-V/2.7 kPa (**Fig. 7e,f**). FA area was smaller on Col-I/13 kPa than on Col-I/2.7 kPa, converging toward the compact adhesions observed on Col-V/2.7 kPa (**Fig. 7g,h**; compare **Fig. 1j**). Thus, raising substrate stiffness roughly fivefold on Col-I reproduced the traction level, spread area, and adhesion morphology that Col-V supports on the soft substrate. These results show that the soft Col-I phenotype does not reflect an intrinsic inability of Col-I to support high traction, and that Col-V presentation produces an adhesion state that, on Col-I, requires a stiffer substrate. Whether Col-V further increases traction on stiff substrates, or whether its effect saturates, was not tested here.

**Figure 7.**
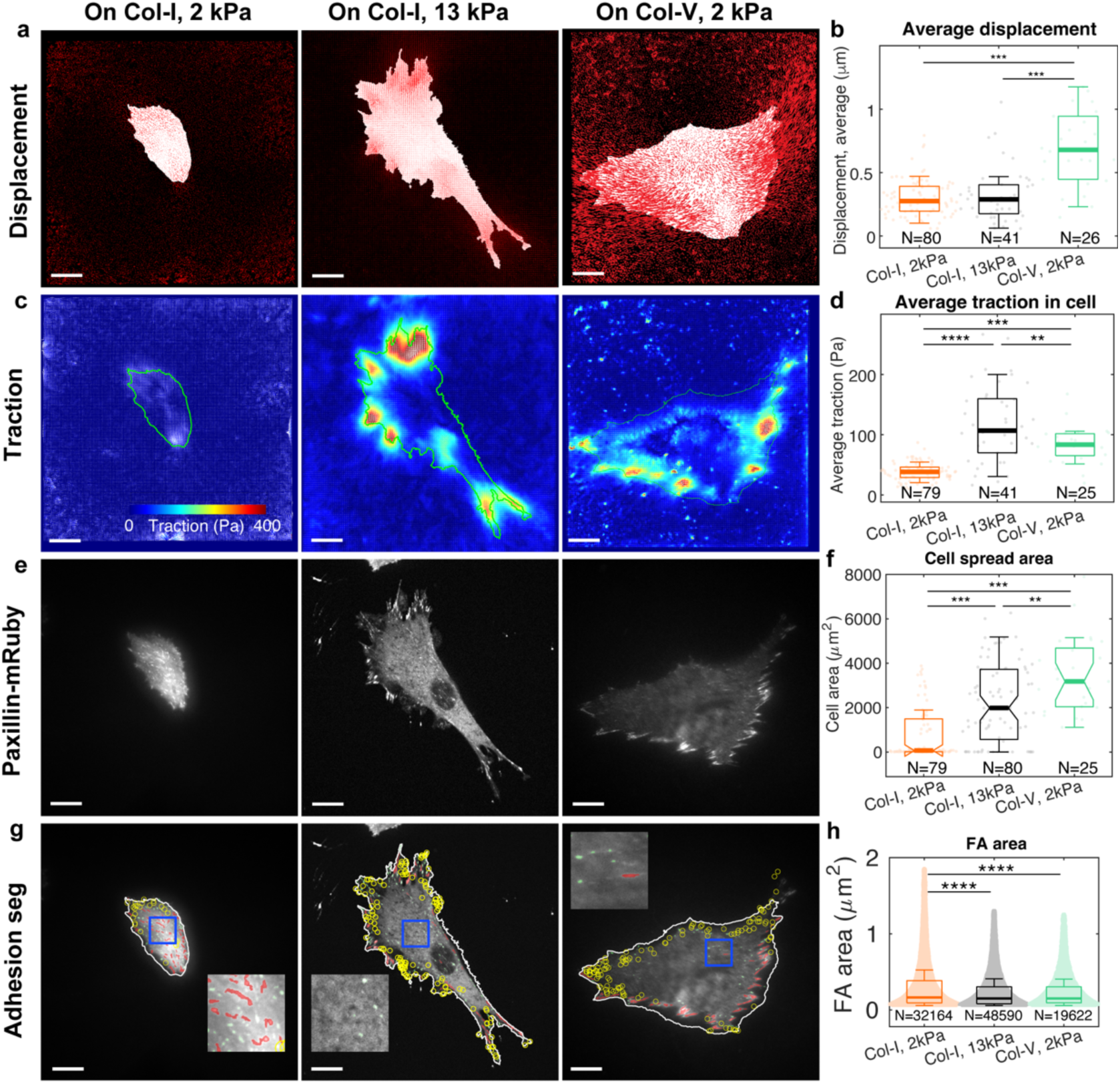
Increasing substrate stiffness reproduces the traction level and compact adhesion phenotype of Col-V on Col-I. Fibroblasts on 2.7 kPa Q-gel substrates coated with Col-I (Col-I/2.7 kPa) or Col-V (Col-V/2.7 kPa) are compared with fibroblasts on 13 kPa Q-gel coated with Col-I (Col-I/13 kPa). **(a)** Representative maps of bead displacement beneath single cells in each condition. **(b)** Average bead displacement per cell. Displacement is comparable between Col-I/2.7 kPa and Col-I/13 kPa and significantly larger on Col-V/2.7 kPa, indicating that raising gel stiffness does not by itself increase substrate deformation. **(c)** Representative traction-stress maps for the same conditions. **(d)** Average traction stress per cell. Because traction scales with the product of local stiffness and displacement, comparable displacements at a roughly fivefold higher stiffness correspond to significantly greater traction on Col-I/13 kPa than on Col-I/2.7 kPa, reaching the level measured on Col-V/2.7 kPa. **(e)** Representative paxillin-mRuby fluorescence images used to outline cells for spread-area measurement. **(f)** Cell spread area, larger on Col-I/13 kPa than on Col-I/2.7 kPa and comparable to Col-V/2.7 kPa. **(g)** Representative focal-adhesion segmentation from paxillin-mRuby TIRF imaging (yellow outlines, with zoomed insets), used to quantify FA area in (h). **(h)** Mean focal-adhesion area, significantly smaller on Col-I/13 kPa than on Col-I/2.7 kPa and converging toward the compact phenotype produced by Col-V on soft gels (compare Fig. 1j). Together, (b–h) show that raising substrate stiffness reproduces both the traction level and the adhesion morphology that Col-V supports on a soft substrate. Box plots show the median and interquartile range with individual cells overlaid; N (number of cells) for each condition is given beneath the corresponding box in (b), (d), and (f). In (h), N denotes the number of individual focal adhesions segmented per condition (pooled across all imaged cells), not the number of cells. Scale bars: 20 µm. Statistical significance (one-way ANOVA with Tukey’s post-hoc test): *P < 0.05, **P < 1 × 10⁻³, ***P < 1 × 10⁻⁵, ****P < 1 × 10⁻⁸.

## 4. Discussion

This study shows that the collagen type presented at the cell-material interface can shift fibroblasts into a mechanical state that otherwise requires a stiffer substrate. On soft silicone substrates of identical stiffness, Col-V presentation increased traction and spreading and produced smaller, more heavily loaded adhesions than Col-I (Fig. 1), accelerated adhesion assembly and disassembly (Fig. 2), and increased migration persistence and collective gap closure (Figs. 3 and 4). Raising substrate stiffness reproduced the traction, spreading, and adhesion morphology of cells on Col-V while keeping Col-I as the ligand (Fig. 7). Separately, in reconstituted hydrogels, Col-V co-assembly produced thinner, straighter fibrils and a stiffer network (Figs. 5 and 6). Below, we discuss what these observations imply for adhesion mechanics and migration, what could make a Col-V interface behave like a stiffer one, and how the hydrogel results bear on the use of Col-V in collagen-based biomaterials.

### 4.1. Focal-adhesion reinforcement and the molecular clutch

The smaller, more heavily loaded adhesions and higher traction per adhesion on Col-V (Fig. 1) can be interpreted within the molecular clutch framework, in which integrin-talin-actin linkages transmit force to the substrate and force-dependent reinforcement, notably through vinculin recruitment, determines whether this coupling is sustained or dissipated [52, 53]. Clutch models predict that traction rises with substrate stiffness up to an optimum, and that changes in clutch properties, such as the number of engaged bonds or their force-dependent lifetimes, can shift this relationship so that cells sustain high traction on compliant substrates [53, 54]. Col-V presentation behaved as such a shift. On a soft substrate, it increased the force borne by each mature adhesion and accelerated both assembly and disassembly (Figs. 1 and 2), signatures of a more dynamically engaged clutch [36, 44, 45]. The stiffness experiment (Fig. 7) supports this interpretation by showing that the same adhesion and traction state is reached on Col-I when substrate resistance is increased. FAs became smaller rather than larger at higher stiffness in our system, in line with multiscale models of fibrous, remodelable matrices that predict smaller adhesions with increasing effective stiffness [55]. Collagen-integrin engagement also differs from fibronectin-integrin engagement in that the collagen-binding motif for integrin α2β1 activates FAK in a tension-independent manner [56], suggesting that the differences we observe arise more from how force is transmitted through adhesions than from FAK-level mechanosensing.

### 4.2. What could make a Col-V interface behave like a stiffer substrate?

Because both coatings share the same bulk substrate, the Col-V effect must arise at the interface. Several non-exclusive mechanisms could contribute. First, Col-I and Col-V are bound by an overlapping set of collagen-binding β1 integrins, but with different structural requirements and relative contributions of α1β1 and α2β1 [17]. A different balance of receptor engagement or bond kinetics on Col-V could change clutch reinforcement without a Col-V-specific receptor. Second, the density, orientation, and conformation of covalently coupled molecules may differ between the two collagens, and ligand density is known to modulate cell responses to substrate compliance [32, 33]. Third, the coupling chemistry may contribute. EDC reacts with collagen carboxyl groups, including those of the glutamate residues in integrin-binding GFOGER-type motifs, and EDC crosslinking reduces integrin-dependent adhesion to collagen in a concentration-dependent manner [57, 58]. Although we used a low EDC concentration for a brief coupling step and treated both collagens identically, the two molecules may differ in how many integrin sites are consumed, which could contribute to the weaker adhesion phenotype on Col-I. However, the wound-closure assay used plasma-treated plastic without EDC, and Col-V still accelerated closure, indicating that the coupling chemistry alone does not account for the Col-V effect. Fourth, both collagens used here were bovine and solubilized by limited pepsin digestion, so neither is expected to retain intact telopeptides, and the Col-V preparation is likely to lack part of the N-terminal noncollagenous domain that projects from heterotypic fibrils in tissue [9], so the effects we report should not be attributed to that domain. Distinguishing among these mechanisms will require characterizing coating density and conformation and comparing coupling chemistries.

### 4.3. A biphasic, adhesion-strength view of the speed-persistence trade-off

The reduced instantaneous speed but increased directional persistence of fibroblasts on Col-V (**Fig. 3**) at first appears counterintuitive, but it aligns closely with the classical biphasic relationship between adhesion strength and migration speed [59]: migration is fastest at intermediate adhesion strength, where traction generated at the front is balanced by efficient adhesion release at the rear, whereas both very low and very high adhesion strength slow migration, the former from insufficient traction and the latter from impaired rear release [59, 60]. The smaller but far more heavily loaded FAs we observe on Col-V (**Fig. 1j**), reflecting higher force per unit adhesion area, are consistent with a shift toward the high-adhesion-strength regime of this model, in which rear-release becomes rate-limiting for instantaneous speed, while the stronger front-rear mechanical coupling stabilizes the axis of movement and increases persistence. This interpretation also offers a point of contrast with the more recently described universal coupling between speed and persistence (UCSP), in which filamentous actin retrograde flow advects polarity cues to reinforce positive feedback between higher speed and higher persistence [61]. Our Col-V data depart from this coupling: persistence increases while speed decreases. We suggest that integrin-mediated adhesion reinforcement, rather than F-actin-flow-driven polarity advection, is the dominant determinant of persistence in this system, and that sufficiently strong adhesion reinforcement can decouple persistence from speed. This may represent a useful, testable extension of the UCSP framework to strongly adhesive, slow-migrating mesenchymal contexts.

### 4.4. From single-cell persistence to collective wound closure

The translation of this single-cell phenotype into faster gap closure (Fig. 4) is consistent with the general principle that, in collective migration, coordination and directional consistency across the monolayer, rather than individual cell speed, are primary determinants of sheet advancement [62–64]. A monolayer of individually slower but more persistent cells can close a gap more efficiently than one of faster but more erratic cells, because net forward progress accumulates only when direction is maintained. We did not measure cell-cell coordination directly, so the link between single-cell persistence and collective closure remains an inference.

### 4.5. Col-V as a component of collagen biomaterials: thinner, straighter fibrils and a stiffer network

The combination of decreased fibril diameter (Fig. 5g) and increased hydrogel stiffness (Fig. 6b) appears paradoxical if fibrils are treated as simple elastic beams, since bending rigidity scales steeply with diameter. This paradox is resolved by the mechanics of disordered networks of stiff fibers such as collagen, which accommodate small deformations mainly through fiber bending, a soft, non-affine mode, until the network transitions to a stretching-dominated regime with a much higher modulus [65, 66]. The strain at which this crossover occurs depends not only on fiber diameter but also on network geometry, including whether fibers are straight or slack [66, 67]. The reduced curvature in Col-V-containing networks (Fig. 5h) indicates fibrils closer to a straightened state, which should lower the strain needed to engage the stiffer stretching mode. Col-V therefore need not make individual fibrils stiffer to stiffen the network; reducing fibril slack may be sufficient. This idea extends the established role of Col-V in limiting lateral fibril growth [46, 68] into a testable prediction for nonlinear network elasticity that could be probed by strain-sweep rheology. For biomaterial design, these results suggest that a small fraction of Col-V could be used to tune the architecture and stiffness of collagen hydrogels without changing total collagen concentration, which otherwise couples stiffness to ligand density and pore size. Whether such Col-V-containing hydrogels reproduce, in 3D, the fibroblast phenotypes we observed on Col-V coatings remains to be tested.

### 4.6. Possible relevance to classical Ehlers-Danlos syndrome

Because cEDS results from reduced Col-V, our gain-of-function experiments can only generate hypotheses for the disease. If the interface effect we observed also operates in tissue, a lower Col-V content in the dermal matrix would be expected to reduce fibroblast traction and migration persistence and to slow wound closure, consistent with delayed healing in cEDS patients [1, 2, 23] and with the rescue of cEDS fibroblast migration by exogenous Col-V [12, 13]. Testing this prediction will require primary dermal fibroblasts from patients and matrices with physiologically relevant Col-V content. Our data also bear on an open question, namely whether connective-tissue phenotypes in EDS are explained by collagen ultrastructural abnormalities alone or whether fibroblast mechanics contributes independently [69]. The coating experiments show that Col-V can change fibroblast mechanics without any change in bulk substrate stiffness or fibril architecture, suggesting that the two explanations are not mutually exclusive. These results are consistent with our earlier proposal that Col-V acts as a mechanical modulator of fibroblast behavior [34], while narrowing it: the present data support an effect at the cell-matrix interface, and the contribution of fibril architecture to fibroblast behavior remains to be established in cells.

The gain-of-function direction may also be informative beyond cEDS. Elevated COL5A1 expression is a recurrent feature of tumor stroma and is associated with disease progression and poor prognosis in several cancers [70]. If stromal Col-V similarly increases fibroblast traction and migration persistence, it could contribute to stromal remodeling and invasion, a possibility that could be tested in cancer-associated fibroblast models.

### 4.7. Limitations

Several aspects of our design limit interpretation. First, the cell-level experiments used thin collagen coatings (covalently coupled for TFM, adhesion imaging, and single-cell migration, and adsorbed on plasma-treated plastic for the wound-closure assay), whereas the structural and mechanical measurements used reconstituted 3D hydrogels. We did not characterize the coatings directly (surface density, molecular conformation, or whether any fibrillar assembly occurs at 1 to 5 µg/mL), so we do not claim that the fibril and network features in Figs. 5 and 6 are present on the coated surfaces. Second, Col-V was a commercial pepsin-solubilized bovine preparation whose chain composition and N-terminal domain status we did not verify, and Col-V isoforms or processing states may differ in their effects. Third, EDC coupling may modify integrin-binding carboxylates [57, 58], and this modification may not affect Col-I and Col-V equally, although the Col-V advantage persisted in the EDC-free wound-closure assay. Fourth, we did not test Col-V on the stiffer substrate, so we cannot say whether Col-V and stiffness act additively or converge on a common limit. Fifth, the Col-V fractions in our hydrogels (10% and 20% by mass) were chosen to produce measurable structural effects; they approach the upper range reported for corneal stroma and exceed the few percent typical of dermis [5, 46]. Sixth, our cellular assays used NIH/3T3 fibroblasts, an immortalized mouse line, cultured in 2D without ascorbate supplementation. TFM and adhesion imaging were performed within 5 h of seeding, so cell-derived matrix is unlikely to have influenced those measurements, but its contribution to the longer migration and wound-closure assays was not measured. Primary human dermal fibroblasts, including cells from cEDS patients, and 3D matrices will be needed to establish physiological relevance. Finally, our stiffness measurements are network-level, and because Col-V content, fibril diameter, and fibril curvature co-vary in our hydrogels, we cannot attribute network stiffening to reduced curvature independently of the change in diameter.

## 5. Conclusion

Presenting Col-V instead of Col-I on a soft substrate shifted fibroblasts into a state of higher traction, smaller and more heavily loaded adhesions, faster adhesion turnover, more persistent migration, and faster collective gap closure, a state that on Col-I required a roughly fivefold stiffer substrate. In reconstituted hydrogels, Col-V co-assembly produced thinner, straighter fibrils and a stiffer network. Together, these findings identify the collagen type presented at the cell-material interface as a design parameter that can partly substitute for bulk stiffness, and they suggest Col-V as a tunable component of collagen-based coatings and scaffolds intended to promote fibroblast migration and repair. Whether loss of this interface effect contributes to delayed wound healing in cEDS remains to be tested.

## Acknowledgements

We thank Smitha Rao and Victoria Santillan (Michigan Technological University) for providing access to the EVOS imaging system used in the wound-closure assay, and Erico Freitas (Materials Characterization and Fabrication Core Facility, Michigan Technological University) for his assistance with scanning electron microscopy. We are especially grateful to Henry Wallace and the Wallace Research Foundation for their generous and continued support of this research. The contributions of the NIH authors were made as part of their official duties as NIH federal employees, are in compliance with agency policy requirements, and are considered Works of the United States Government. The findings and conclusions presented in this paper are those of the authors and do not necessarily reflect the views of the NIH or the US Department of Health and Human Services.

## Funding

This work was supported by the Wallace Research Foundation (WALLA000, WALLA001) and the National Institutes of Health (1R15GM135806) to H.T, S.R., M.R., and S.J.H. This work was supported in part by the Intramural Research Program of the National Institutes of Health (NIH), National Institute of Biomedical Imaging and Bioengineering with grant number ZIA-EB000094 and the NIH’s Distinguished Scholars Program to A.X.C.R and D.C.

## CRediT authorship contribution statement

**Shaina P. Royer:** Conceptualization, Investigation, Methodology, Formal analysis, Writing – original draft. **Himja Tiwari:** Methodology, Formal analysis. **David Cho:** Methodology, Formal analysis. **Sesha Sai Ramineni:** Methodology, Formal analysis. **Majid Rostami:** Methodology, Formal analysis. **Mohanish Chandurkar:** Investigation, Methodology. **Alexander X. Cartagena-Rivera:** Methodology, Formal analysis, Funding acquisition, Writing – review & editing. **Sangyoon J. Han:** Conceptualization, Supervision, Funding acquisition, Writing – original draft, Writing – review & editing.

## Declaration of competing interest

The authors declare that they have no known competing financial interests or personal relationships that could have appeared to influence the work reported in this paper.

## Data availability

All data supporting the findings of this study are available from the corresponding author upon reasonable request. The Focal Adhesion Package software used for adhesion detection, tracking, and classification is publicly available at https://github.com/HanLab-BME-MTU/focalAdhesionPackage.git.

## Notes

### Competing Interest Statement

The authors have declared no competing interest.

## References

[1] F. Malfait, C. Francomano, P. Byers, J. Belmont, B. Berglund, J. Black, L. Bloom, J.M. Bowen, A.F. Brady, N.P. Burrows, M. Castori, H. Cohen, M. Colombi, S. Demirdas, J. De Backer, A. De Paepe, S. Fournel-Gigleux, M. Frank, N. Ghali, C. Giunta, R. Grahame, A. Hakim, X. Jeunemaitre, D. Johnson, B. Juul-Kristensen, I. Kapferer-Seebacher, H. Kazkaz, T. Kosho, M.E. Lavallee, H. Levy, R. Mendoza-Londono, M. Pepin, F.M. Pope, E. Reinstein, L. Robert, M. Rohrbach, L. Sanders, G.J. Sobey, T. Van Damme, A. Vandersteen, C. van Mourik, N. Voermans, N. Wheeldon, J. Zschocke, B. Tinkle, The 2017 international classification of the Ehlers-Danlos syndromes, Am J Med Genet C Semin Med Genet 175(1) (2017) 8–26.

[2] J.M. Bowen, G.J. Sobey, N.P. Burrows, M. Colombi, M.E. Lavallee, F. Malfait, C.A. Francomano, Ehlers-Danlos syndrome, classical type, Am J Med Genet C Semin Med Genet 175(1) (2017) 27–39.

[3] S. Symoens, D. Syx, F. Malfait, B. Callewaert, J. De Backer, O. Vanakker, P. Coucke, A. De Paepe, Comprehensive molecular analysis demonstrates type V collagen mutations in over 90% of patients with classic EDS and allows to refine diagnostic criteria, Hum Mutat 33(10) (2012) 1485–1493.

[4] R.P. Boot-Handford, D.S. Tuckwell, Fibrillar collagen: the key to vertebrate evolution? A tale of molecular incest, Bioessays 25(2) (2003) 142–51.

[5] K.M. Mak, C.Y. Png, D.J. Lee, Type V Collagen in Health, Disease, and Fibrosis, Anat Rec (Hoboken) 299(5) (2016) 613–29.

[6] C. Catala-Petavy, L. Machet, G. Georgesco, F. Petavy, A. Maruani, L. Vaillant, Contribution of skin biometrology to the diagnosis of the Ehlers-Danlos syndrome in a prospective series of 41 patients, Skin Res Technol 15(4) (2009) 412–7.

[7] M. Neshatian, N. Mittal, S. Huang, A. Ali, E. Khattignavong, L. Bozec, Investigation of dermal collagen nanostructures in Ehlers-Danlos Syndrome (EDS) patients, PLoS One 19(8) (2024) e0307442.

[8] I. Hausser, I. Anton-Lamprecht, Differential ultrastructural aberrations of collagen fibrils in Ehlers-Danlos syndrome types I-IV as a means of diagnostics and classification, Hum Genet 93(4) (1994) 394–407.

[9] T.F. Linsenmayer, E. Gibney, F. Igoe, M.K. Gordon, J.M. Fitch, L.I. Fessler, D.E. Birk, Type V collagen: molecular structure and fibrillar organization of the chicken alpha 1(V) NH2-terminal domain, a putative regulator of corneal fibrillogenesis, J Cell Biol 121(5) (1993) 1181–9.

[10] D.E. Birk, J.M. Fitch, J.P. Babiarz, K.J. Doane, T.F. Linsenmayer, Collagen fibrillogenesis in vitro: interaction of types I and V collagen regulates fibril diameter, J Cell Sci 95(Pt 4) (1990) 649–657.

[11] R.J. Wenstrup, J.B. Florer, J.M. Davidson, C.L. Phillips, B.J. Pfeiffer, D.W. Menezes, I. Chervoneva, D.E. Birk, Murine model of the Ehlers-Danlos syndrome. col5a1 haploinsufficiency disrupts collagen fibril assembly at multiple stages, J Biol Chem 281(18) (2006) 12888–12895.

[12] S. Viglio, N. Zoppi, A. Sangalli, A. Gallanti, S. Barlati, M. Mottes, M. Colombi, M. Valli, Rescue of migratory defects of Ehlers-Danlos syndrome fibroblasts in vitro by type V collagen but not insulin-like binding protein-1, J Invest Dermatol 128(8) (2008) 1915–9.

[13] A.D. Berendsen, A.L. Bronckers, T.H. Smit, X.F. Walboomers, V. Everts, Collagen type V enhances matrix contraction by human periodontal ligament fibroblasts seeded in three-dimensional collagen gels, Matrix Biol 25(8) (2006) 515–22.

[14] X. Pang, X. He, Z. Qiu, H. Zhang, R. Xie, Z. Liu, Y. Gu, N. Zhao, Q. Xiang, Y. Cui, Targeting integrin pathways: mechanisms and advances in therapy, Signal Transduct Target Ther 8(1) (2023) 1.

[15] C.G. Knight, L.F. Morton, A.R. Peachey, D.S. Tuckwell, R.W. Farndale, M.J. Barnes, The collagen-binding A-domains of integrins alpha(1)beta(1) and alpha(2)beta(1) recognize the same specific amino acid sequence, GFOGER, in native (triple-helical) collagens, J Biol Chem 275(1) (2000) 35–40.

[16] M. Musiime, J. Chang, U. Hansen, K.E. Kadler, C. Zeltz, D. Gullberg, Collagen Assembly at the Cell Surface: Dogmas Revisited, Cells 10(3) (2021).

[17] F. Ruggiero, J. Comte, C. Cabanas, R. Garrone, Structural requirements for alpha 1 beta 1 and alpha 2 beta 1 integrin mediated cell adhesion to collagen V, J Cell Sci 109 (Pt 7) (1996) 1865–74.

[18] N. Lu, T.V. Karlsen, R.K. Reed, M. Kusche-Gullberg, D. Gullberg, Fibroblast alpha11beta1 integrin regulates tensional homeostasis in fibroblast/A549 carcinoma heterospheroids, PLoS One 9(7) (2014) e103173.

[19] E.J. Hunter, S.W. Hamaia, D. Gullberg, J.D. Malcor, R.W. Farndale, Selectivity of the collagen-binding integrin inhibitors, TC-I-15 and obtustatin, Toxicol Appl Pharmacol 428 (2021) 115669.

[20] M.J. Humphries, The molecular basis and specificity of integrin-ligand interactions, J Cell Sci 97 (Pt 4) (1990) 585–92.

[21] J.D. Humphries, A. Byron, M.J. Humphries, Integrin ligands at a glance, J Cell Sci 119(Pt 19) (2006) 3901–3.

[22] N. Mittal, E.B. Michels, A.E. Massey, Y. Qiu, S.P. Royer-Weeden, B.R. Smith, A.X. Cartagena-Rivera, S.J. Han, Myosin-independent stiffness sensing by fibroblasts is regulated by the viscoelasticity of flowing actin, Communications Materials 5(1) (2024) 6.

[23] J. DeNigris, Q. Yao, E.K. Birk, D.E. Birk, Altered dermal fibroblast behavior in a collagen V haploinsufficient murine model of classic Ehlers-Danlos syndrome, Connect Tissue Res 57(1) (2016) 1–9.

[24] H. Levinson, A Paradigm of Fibroblast Activation and Dermal Wound Contraction to Guide the Development of Therapies for Chronic Wounds and Pathologic Scars, Adv Wound Care (New Rochelle) 2(4) (2013) 149–159.

[25] D.I. Shreiber, V.H. Barocas, R.T. Tranquillo, Temporal variations in cell migration and traction during fibroblast-mediated gel compaction, Biophys J 84(6) (2003) 4102–14.

[26] S.J. Han, M.L. Rodriguez, Z. Al-Rekabi, N.J. Sniadecki, Spatial and Temporal Coordination of Traction Forces in One-Dimensional Cell Migration, Cell Adhesion & Migration (2016) 00–00.

[27] J.T. Parsons, A.R. Horwitz, M.A. Schwartz, Cell adhesion: integrating cytoskeletal dynamics and cellular tension, Nat Rev Mol Cell Biol 11(9) (2010) 633–43.

[28] M. D’Urso, N.A. Kurniawan, Mechanical and Physical Regulation of Fibroblast-Myofibroblast Transition: From Cellular Mechanoresponse to Tissue Pathology, Front Bioeng Biotechnol 8 (2020) 609653.

[29] C. Lo, H. Wang, M. Dembo, Y. Wang, Cell movement is guided by the rigidity of the substrate, Biophys J 79(1) (2000) 144–152.

[30] R.J. Pelham, Jr., Y. Wang, Cell locomotion and focal adhesions are regulated by substrate flexibility, Proc Natl Acad Sci U S A 94(25) (1997) 13661–5.

[31] S. Asano, S. Ito, K. Takahashi, K. Furuya, M. Kondo, M. Sokabe, Y. Hasegawa, Matrix stiffness regulates migration of human lung fibroblasts, Physiol Rep 5(9) (2017).

[32] S.J. Han, K.S. Bielawski, L.H. Ting, M.L. Rodriguez, N.J. Sniadecki, Decoupling substrate stiffness, spread area, and micropost density: a close spatial relationship between traction forces and focal adhesions, Biophys J 103(4) (2012) 640–8.

[33] A. Engler, L. Bacakova, C. Newman, A. Hategan, M. Griffin, D. Discher, Substrate compliance versus ligand density in cell on gel responses, Biophys J 86(1 Pt 1) (2004) 617–28.

[34] S.P. Royer, S.J. Han, Mechanobiology in the Comorbidities of Ehlers Danlos Syndrome, Front Cell Dev Biol 10 (2022) 874840.

[35] N. Mittal, H.S. Lee, M.H. Ginsberg, S.J. Han, Regulated RIAM-Talin Engagement Controls Adhesion Stability and Mechanical Output, bioRxiv (2026).

[36] S.J. Han, E.V. Azarova, A.J. Whitewood, A. Bachir, E. Guttierrez, A. Groisman, A.R. Horwitz, B.T. Goult, K.M. Dean, G. Danuser, Pre-complexation of talin and vinculin without tension is required for efficient nascent adhesion maturation, Elife 10 (2021).

[37] M. Rostami, S.J. Han, High-Throughput Focal Adhesion Analysis Software for Automated Detection, Tracking, and Classification of Cell-Matrix Adhesions, Current Protocols (2026).

[38] D.J. Webb, K. Donais, L.A. Whitmore, S.M. Thomas, C.E. Turner, J.T. Parsons, A.F. Horwitz, FAK-Src signalling through paxillin, ERK and MLCK regulates adhesion disassembly, Nat Cell Biol 6(2) (2004) 154–61.

[39] S.J. Han, Y. Oak, A. Groisman, G. Danuser, Traction microscopy to identify force modulation in subresolution adhesions, Nat Methods 12(7) (2015) 653–6.

[40] N. Costigliola, L. Ding, C.J. Burckhardt, S.J. Han, E. Gutierrez, A. Mota, A. Groisman, T.J. Mitchison, G. Danuser, Vimentin fibers orient traction stress, Proc Natl Acad Sci U S A 114(20) (2017) 5195–5200.

[41] A.D. Doyle, Generation of 3D Collagen Gels with Controlled Diverse Architectures, Curr Protoc Cell Biol 72 (2016) 10 20 1–10 20 16.

[42] E.K. Dimitriadis, F. Horkay, J. Maresca, B. Kachar, R.S. Chadwick, Determination of elastic moduli of thin layers of soft material using the atomic force microscope, Biophys J 82(5) (2002) 2798–810.

[43] D.H. Cho, S. Aguayo, A.X. Cartagena-Rivera, Atomic force microscopy-mediated mechanobiological profiling of complex human tissues, Biomaterials 303 (2023) 122389.

[44] M. Nagano, D. Hoshino, N. Koshikawa, T. Akizawa, M. Seiki, Turnover of focal adhesions and cancer cell migration, Int J Cell Biol 2012 (2012) 310616.

[45] M. Mavrakis, M.A. Juanes, The compass to follow: Focal adhesion turnover, Curr Opin Cell Biol 80 (2023) 102152.

[46] D.E. Birk, Type V collagen: heterotypic type I/V collagen interactions in the regulation of fibril assembly, Micron 32(3) (2001) 223–37.

[47] R.J. Wenstrup, J.B. Florer, E.W. Brunskill, S.M. Bell, I. Chervoneva, D.E. Birk, Type V collagen controls the initiation of collagen fibril assembly, J Biol Chem 279(51) (2004) 53331–7.

[48] M.D. Shoulders, R.T. Raines, Collagen structure and stability, Annu Rev Biochem 78 (2009) 929–58.

[49] W.W. Leow, W. Hwang, Epitaxially guided assembly of collagen layers on mica surfaces, Langmuir 27(17) (2011) 10907–13.

[50] F. Jiang, K. Khairy, K. Poole, J. Howard, D.J. Muller, Creating nanoscopic collagen matrices using atomic force microscopy, Microsc Res Tech 64(5-6) (2004) 435–40.

[51] P.D. Garcia, R. Garcia, Determination of the Elastic Moduli of a Single Cell Cultured on a Rigid Support by Force Microscopy, Biophys J 114(12) (2018) 2923–2932.

[52] A. Elosegui-Artola, R. Oria, Y. Chen, A. Kosmalska, C. Perez-Gonzalez, N. Castro, C. Zhu, X. Trepat, P. Roca-Cusachs, Mechanical regulation of a molecular clutch defines force transmission and transduction in response to matrix rigidity, Nat Cell Biol 18(5) (2016) 540–8.

[53] B.L. Bangasser, S.S. Rosenfeld, D.J. Odde, Determinants of maximal force transmission in a motor-clutch model of cell traction in a compliant microenvironment, Biophys J 105(3) (2013) 581–92.

[54] C.E. Chan, D.J. Odde, Traction dynamics of filopodia on compliant substrates, Science 322(5908) (2008) 1687–91.

[55] X. Cao, E. Ban, B.M. Baker, Y. Lin, J.A. Burdick, C.S. Chen, V.B. Shenoy, Multiscale model predicts increasing focal adhesion size with decreasing stiffness in fibrous matrices, Proc Natl Acad Sci U S A 114(23) (2017) E4549–E4555.

[56] J. Seong, A. Tajik, J. Sun, J.L. Guan, M.J. Humphries, S.E. Craig, A. Shekaran, A.J. Garcia, S. Lu, M.Z. Lin, N. Wang, Y. Wang, Distinct biophysical mechanisms of focal adhesion kinase mechanoactivation by different extracellular matrix proteins, Proc Natl Acad Sci U S A 110(48) (2013) 19372–7.

[57] D.V. Bax, N. Davidenko, D. Gullberg, S.W. Hamaia, R.W. Farndale, S.M. Best, R.E. Cameron, Fundamental insight into the effect of carbodiimide crosslinking on cellular recognition of collagen-based scaffolds, Acta Biomater 49 (2017) 218–234.

[58] D.V. Bax, N. Davidenko, S.W. Hamaia, R.W. Farndale, S.M. Best, R.E. Cameron, Impact of UV- and carbodiimide-based crosslinking on the integrin-binding properties of collagen-based materials, Acta Biomater 100 (2019) 280–291.

[59] P.A. DiMilla, K. Barbee, D.A. Lauffenburger, Mathematical model for the effects of adhesion and mechanics on cell migration speed, Biophys J 60(1) (1991) 15–37.

[60] S.P. Palecek, J.C. Loftus, M.H. Ginsberg, D.A. Lauffenburger, A.F. Horwitz, Integrin-ligand binding properties govern cell migration speed through cell-substratum adhesiveness, Nature 385(6616) (1997) 537–40.

[61] P. Maiuri, J.F. Rupprecht, S. Wieser, V. Ruprecht, O. Benichou, N. Carpi, M. Coppey, S. De Beco, N. Gov, C.P. Heisenberg, C. Lage Crespo, F. Lautenschlaeger, M. Le Berre, A.M. Lennon-Dumenil, M. Raab, H.R. Thiam, M. Piel, M. Sixt, R. Voituriez, Actin flows mediate a universal coupling between cell speed and cell persistence, Cell 161(2) (2015) 374–86.

[62] P. Vitorino, T. Meyer, Modular control of endothelial sheet migration, Genes Dev 22(23) (2008) 3268–81.

[63] A. Zaritsky, D. Kaplan, I. Hecht, S. Natan, L. Wolf, N.S. Gov, E. Ben-Jacob, I. Tsarfaty, Propagating waves of directionality and coordination orchestrate collective cell migration, PLoS Comput Biol 10(7) (2014) e1003747.

[64] P. Friedl, D. Gilmour, Collective cell migration in morphogenesis, regeneration and cancer, Nat Rev Mol Cell Biol 10(7) (2009) 445–57.

[65] C. Heussinger, B. Schaefer, E. Frey, Nonaffine rubber elasticity for stiff polymer networks, Phys Rev E Stat Nonlin Soft Matter Phys 76(3 Pt 1) (2007) 031906.

[66] A.J. Licup, S. Munster, A. Sharma, M. Sheinman, L.M. Jawerth, B. Fabry, D.A. Weitz, F.C. MacKintosh, Stress controls the mechanics of collagen networks, Proc Natl Acad Sci U S A 112(31) (2015) 9573–8.

[67] C.P. Broedersz, F.C. MacKintosh, Modeling semiflexible polymer networks, Reviews of Modern Physics 86(3) (2014) 995–1036.

[68] E.G. Canty, K.E. Kadler, Collagen fibril biosynthesis in tendon: a review and recent insights, Comp Biochem Physiol A Mol Integr Physiol 133(4) (2002) 979–85.

[69] S. Malek, D.V. Koster, The Role of Cell Adhesion and Cytoskeleton Dynamics in the Pathogenesis of the Ehlers-Danlos Syndromes and Hypermobility Spectrum Disorders, Front Cell Dev Biol 9 (2021) 649082.

[70] X. Chen, C. Ma, Y. Li, Y. Liang, T. Chen, D. Han, D. Luo, N. Zhang, W. Zhao, L. Wang, Q. Yang, COL5A1 promotes triple-negative breast cancer progression by activating tumor cell-macrophage crosstalk, Oncogene 43(23) (2024) 1742–1756.

