## Supplemental Information for "Collagen V Presented at the Cell-Substrate Interface Mimics Substrate Stiffening to Enhance Fibroblast Traction, Adhesion Turnover, and Persistent Migration"

**Supplementary Note 1. Integrin specificity of Col-V**

A distinct, Col-V–specific integrin heterodimer might in principle explain Col-V's unique biological effects; however, the literature argues strongly against this possibility. Fibrillar collagens are recognized predominantly by the “collagen-binding” $\beta_1$-integrins, *i.e.*, $\alpha_1\beta_1$, $\alpha_2\beta_1$, $\alpha_{10}\beta_1$, and $\alpha_{11}\beta_1$, which bind GFOGER-like motifs present across collagens I, II, III, V, and XI [1, 2]. Early adhesion assays demonstrated that both $\alpha_1\beta_1$ and $\alpha_2\beta_1$ mediate cell binding to Col-V triple-helical fragments, with nearly identical sequence requirements to those defined for Col-I [3-6]. Quantitative studies further show that $\alpha_2\beta_1$ binds collagens I–III with high affinity and can also engage Col-V and Col-XI, whereas $\alpha_1\beta_1$ favors network collagens such as Col-IV yet still supports adhesion to Col-V [2]. No integrin has been described that selectively recognizes Col-V over Col-I. Likewise, fibroblast-specific $\alpha_{11}\beta_1$, primarily a Col-I/III receptor, has not been reported as Col-V-specific [7]. This lack of a clearly Col-V–specific receptor strengthens the rationale for focusing on biophysical mechanisms, such as fibril nano-architecture, bending rigidity and gel-level stiffness, rather than novel biochemical receptors to explain how Col-V selectively tunes fibroblast mechanosensing.

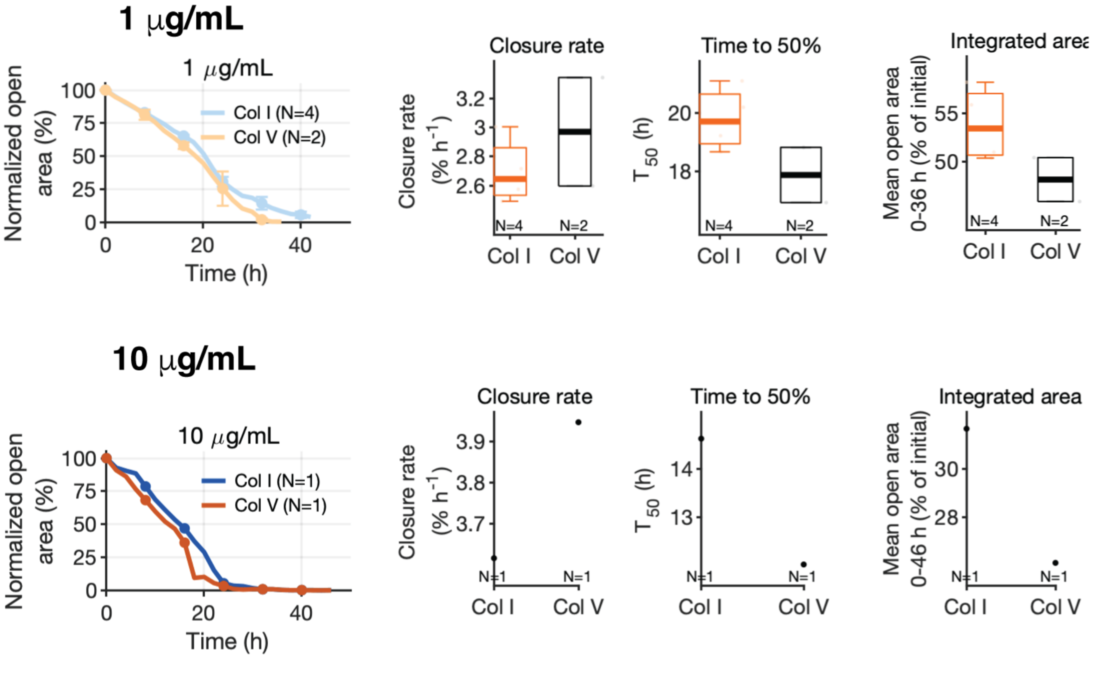


*Supplementary Figure 1. **Wound closure on Col-I and Col-V at 1 and 10 μg/mL coating density.***

(a–d) Wound-closure assay on tissue culture plastic coated with 1 μg/mL Col-I or Col-V. (a) Normalized open-area curves (mean ± SEM), each well normalized to its own open area at t = 0. (b) Wound-closure rate. (c) Time to 50% closure ($T_{50}$). (d) Time-averaged open area over 0–36 hr, the longest interval common to every well at this coating density. (e–h) The same four measurements at 10 μg/mL, with the integration window in (h) running 0–46 hr. All metrics were computed exactly as in Fig. 4, and the box plots, outlier handling and statistical conventions follow that figure. At 1 µg/mL the closure rate (2.97 ± 0.53 vs. 2.70 ± 0.23 % $hr^{-1}$), $T_{50}$ (17.9 ± 1.4 vs. 19.8 ± 1.1 hr) and time-averaged open area (48.1 ± 3.2 vs. 53.9 ± 3.8 %) all favour Col-V on average (mean ± SD, Col-V vs. Col-I), but the distributions overlap and none of the comparisons reaches significance (P = 0.53, 0.27 and 0.27 respectively; N = 4 Col-I, N = 2 Col-V). With these group sizes the smallest attainable two-sided rank-sum P value is 0.13, so the 1 µg/mL comparison is under-powered by construction and should not be read as evidence that no difference exists. At 10 µg/mL only a single well per condition was imaged; those panels are shown descriptively, without error bars or statistical testing, and indicate only that closure is faster on both substrates at this coating density ($T_{50}$ = 12.0 hr on Col-V vs. 14.6 hr on Col-I). Because the 1 µg/mL wells and most of the 5 µg/mL wells were imaged in separate experimental batches at different passage numbers, coating density and batch are partially confounded across this series; the comparison between densities is therefore descriptive rather than a controlled dose–response.

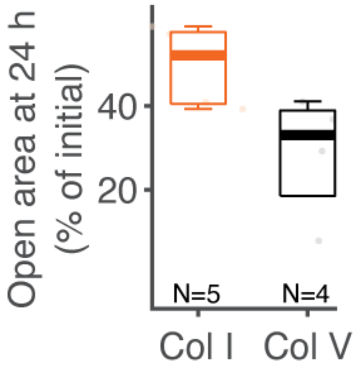


*Supplementary Figure 2.* ***Residual open area at 24 hr for the 5 µg/mL wound-closure assay.***

Residual open area at 24 hr, expressed as a percentage of each well's initial gap. Because 24 hr falls close to the mean $T_{50}$ on Col-I (23.6 hr), that is, on the steepest part of the Col-I curve, a single-frame reading at this time is largely a noisier restatement of $T_{50}$; the main text therefore reports the time-averaged open area instead. The difference here runs in the same direction as every other metric (28.7 ± 14.8 % on Col-V vs. 49.6 ± 9.1 % on Col-I, mean ± SD) but does not reach significance at this replicate number (P = 0.064, two-sided Wilcoxon rank-sum test; N = 5 Col-I, N = 4 Col-V). Box plots, outlier handling and statistics follow the conventions given for Fig. 4.

### References

[1] X. Pang, X. He, Z. Qiu, H. Zhang, R. Xie, Z. Liu, Y. Gu, N. Zhao, Q. Xiang, Y. Cui, Targeting integrin pathways: mechanisms and advances in therapy, Signal Transduct Target Ther 8(1) (2023) 1.

[2] C.G. Knight, L.F. Morton, A.R. Peachey, D.S. Tuckwell, R.W. Farndale, M.J. Barnes, The collagen-binding A-domains of integrins alpha(1)beta(1) and alpha(2)beta(1) recognize the same specific amino acid sequence, GFOGER, in native (triple-helical) collagens, J Biol Chem 275(1) (2000) 35-40.
[3] M. Musiime, J. Chang, U. Hansen, K.E. Kadler, C. Zeltz, D. Gullberg, Collagen Assembly at the Cell Surface: Dogmas Revisited, Cells 10(3) (2021).
[4] J.D. Humphries, A. Byron, M.J. Humphries, Integrin ligands at a glance, J Cell Sci 119(Pt 19) (2006) 3901-3.
[5] F. Ruggiero, J. Comte, C. Cabanas, R. Garrone, Structural requirements for alpha 1 beta 1 and alpha 2 beta 1 integrin mediated cell adhesion to collagen V, J Cell Sci 109 ( Pt 7) (1996) 1865-74.
[6] E.J. Hunter, S.W. Hamaia, D. Gullberg, J.D. Malcor, R.W. Farndale, Selectivity of the collagen-binding integrin inhibitors, TC-I-15 and obtustatin, Toxicol Appl Pharmacol 428 (2021) 115669.
[7] N. Lu, T.V. Karlsen, R.K. Reed, M. Kusche-Gullberg, D. Gullberg, Fibroblast alpha11beta1 integrin regulates tensional homeostasis in fibroblast/A549 carcinoma heterospheroids, PLoS One 9(7) (2014) e103173.

**Supplementary Video Legends**

**Supplementary Video 1. Focal adhesion dynamics of a fibroblast on collagen I.** Time-lapse TIRF microscopy of a paxillin-mRuby–expressing NIH/3T3 fibroblast on a Col-I–coated glass substrate, imaged every 15 seconds for 15 minutes starting 1.5 h after seeding. Individual adhesion trajectories were subsequently color-coded by lifetime status (nascent adhesion, focal complex, or focal adhesion) as shown in Fig. 2. Related to Fig. 2.

**Supplementary Video 2. Focal adhesion dynamics of a fibroblast on collagen V.** Time-lapse TIRF microscopy of a paxillin-mRuby–expressing NIH/3T3 fibroblast on a Col-V–coated glass substrate, imaged and processed as in Supplementary Video 1. Adhesions on Col-V assemble and disassemble more rapidly than on Col-I. Related to Fig. 2.

**Supplementary Video 3. Collective wound closure on collagen I.** Time-lapse phase-contrast imaging of an NIH/3T3 fibroblast monolayer migrating into a cell-free gap defined by a Culture-Insert 2 Well (ibidi) on tissue culture plastic coated with 5 µg/mL Col-I, imaged with a 4× objective every 2 hours for up to 48 hours after insert removal. The gap closes slowly and remains about half open at 24 hr. Related to Fig. 4.

**Supplementary Video 4. Collective wound closure on collagen V.** Time-lapse phase-contrast imaging of an NIH/3T3 fibroblast monolayer migrating into a cell-free gap on tissue culture plastic coated with 5 µg/mL Col-V, imaged and processed as in Supplementary Video 3. Closure is markedly faster: the gap is about one third open at 24 hr and is essentially complete by the end of the imaging window. Related to Fig. 4.
